# Situating the parietal memory network in the context of multiple parallel distributed networks using high-resolution functional connectivity

**DOI:** 10.1101/2023.08.16.553585

**Authors:** Y. Kwon, J.J. Salvo, N. Anderson, A.M Holubecki, M. Lakshman, K. Yoo, K. Kay, C. Gratton, R.M. Braga

**Affiliations:** Northwestern University Department of Neurology; Yale University Department of Psychology; University of Minnesota Department of Radiology; Florida State University Department of Psychology

## Abstract

A principle of brain organization is that networks serving higher cognitive functions are widely distributed across the brain. One exception has been the parietal memory network (PMN), which plays a role in recognition memory but is often defined as being restricted to posteromedial association cortex. We hypothesized that high-resolution estimates of the PMN would reveal small regions that had been missed by prior approaches. High-field 7T functional magnetic resonance imaging (fMRI) data from extensively sampled participants was used to define the PMN within individuals. The PMN consistently extended beyond the core posteromedial set to include regions in the inferior parietal lobule; rostral, dorsal, medial, and ventromedial prefrontal cortex; the anterior insula; and ramus marginalis of the cingulate sulcus. The results suggest that, when fine-scale anatomy is considered, the PMN matches the expected distributed architecture of other association networks, reinforcing that parallel distributed networks are an organizing principle of association cortex.

---

The cerebral cortices comprise large-scale networks that are specialized for different cognitive functions (Doucet et al., 2011; Geschwind, 1965; Goldman-Rakic, 1988; Mesulam, 1981, 1990; Power et al., 2011; Smith et al., 2009; Yeo et al., 2011). Knowledge of the detailed anatomy of the networks, including the constituent regions and how these fit within broader topographical patterns, can provide clues to the component processes of cognition and their anatomical basis (e.g., Kanwisher, 2010; Krubitzer, 2007; Margulies et al., 2016). One example is recognition (Squire, 1986; Tulving 1983), a type of declarative memory that is commonly divided into processes of recollection (i.e., the mental re-experiencing of a previous experience) and familiarity (i.e., the subjective feeling that something has been previously experienced; Cabeza et al., 2008; Johnson et al. 1993; Yonelinas et al., 2002, 2005). Although these processes likely interact during recognition, evidence from functional brain imaging supports that distinct brain systems are associated with recollection and familiarity (e.g., Cabeza et al., 2008; J. Chen et al., 2017; Wheeler & Buckner, 2004).

Tasks targeting recollection often reveal increased activity in a broad network that includes regions within the canonical ‘default network’ or ‘DN’ (Addis et al., 2007; Buckner et al., 2008; DiNicola et al., 2020; Wagner et al., 2005). The implicated network is widely distributed, including regions at or near the posterior cingulate, posterior inferior parietal, lateral temporal, medial and lateral prefrontal, and parahippocampal cortices (Buckner et al., 2008; Vincent et al., 2006). This network shows increased activity when a participant is asked to think about a past or prospective future event (Addis et al., 2007), tracks participant’s subjective ratings of how vividly they mentally experienced the thought-of event (DiNicola et al., 2023), and is robustly activated when mental scenes are contemplated (DiNicola et al., 2023; Peer et al., 2015; Silson et al., 2019). In contrast, tasks that target familiarity, such as those involving detection of previously presented stimuli (i.e., old/new effects), typically reveal activity in a much more restricted posteromedial set of regions, a network sometimes called the ‘parietal memory network’ or ‘PMN’ (Gilmore et al., 2015, 2019). The PMN includes prominent regions at or near the precuneus (PCU) and rostral posterior cingulate cortex (rPCC) within the callosal sulcus. These PMN regions surround the posteromedial regions of the canonical DN (Doucet et al., 2011; Power et al., 2011; Yeo et al., 2011) which forms a key identifying feature of the PMN. Functionally, the PMN shows increased activity to previously seen stimuli – the so-called ‘repetition enhancement effect’ (Kim, 2017; Segaert et al., 2012) – even in the absence of an explicit requirement for the stimuli to be identified as familiar (Gilmore et al., 2019; see also Rosen et al., 2018). Hence, evidence supports that two distinct networks, one within canonical DN regions and one being the PMN, play complementary but dissociable roles in recognition.

The DN is a widely distributed network, containing regions in multiple association areas, in an organizational motif that is thought to be characteristic of association cortex (Goldman-Rakic, 1988; Margulies et al., 2016). In contrast, the PMN is sometimes described as being restricted to posteromedial cortex (i.e., the PCU and rPCC). This bears scrutiny given that the PMN is also located deep within association cortex and appears to serve higher-order associative functions. These observations suggest that the PMN should also have a distributed network organization that more resembles the DN and other association networks (e.g., see Braga & Buckner, 2017; DiNicola & Buckner, 2021). Analysis of blood-oxygenation-level-dependent (BOLD) signal correlations (i.e., functional connectivity or FC; Biswal et al., 1995) is useful for investigating the organization of brain networks, because it provides estimates that do not rely on the specifics of a given task, and yet largely correspond to task-evoked activity patterns across multiple domains (e.g., see Fox et al., 2006; Power et al., 2011; Salvo et al., 2021; Smith et al., 2009; Tavor et al., 2016). Some prior FC estimates have suggested that the rPCC and PCU regions of the PMN are part of a subsystem of the canonical DN (e.g., Barnett et al. 2021; Cooper et al. 2019; 2021; Power et al. 2011), which would accord with the PMN and DN’s similar functions supporting recognition. Power et al. (2011) indicated that the PCU and rPCC regions cluster with the DN at lower dimensionality (see the 1^st^ figure in Power et al., 2011), but separate into a distinct network (i.e., the PMN) at higher dimensionality (see the 4^th^ figure in Power et al., 2011). However, these same PCU and rPCC regions are considered part of the frontoparietal control network (FPN) in the 7-network parcellation of Yeo et al. (2011; see their 7^th^ figure; see also Vincent et al., 2008), which then separates into a distinct network (i.e., the PMN) in their 17-network solution (see the 9th figure of Yeo et al., 2011; see also Dixon et al., 2018). Clarifying how the PMN relates to or is distinct from these nearby and perhaps broader systems could provide clues to the cognitive processes supported by the PMN. The FPN is considered a ‘multiple demand’ system thought to be recruited for multiple tasks when additional cognitive resources are needed (e.g., when difficulty is increased; Duncan, 2010). Hence a closer link between the PMN and FPN may point to the PMN being a domain-general system that is not exclusively recruited for recognition-related processes.

Alternatively, it is possible that the PMN is a distinct system that has been blurred together with nearby networks in prior estimates. Even when the PMN is defined as a distinct network, there have been conflicting accounts of which regions are involved (reviewed in Gilmore et al., 2021). Some FC estimates restrict the PMN to the core PCU and rPCC regions, particularly when winner-takes-all algorithms are used (e.g., compare the 13^th^ and 34^th^ figures in Yeo et al., 2011; see also Doucet et al., 2011; Power et al., 2011; Schaefer et al., 2018), but several studies have identified additional PMN regions. A PMN region is often reported in the inferior parietal lobule (IPL), sometimes at or near the posterior angular gyrus (e.g., Gilmore et al., 2015, 2021; Gordon et al., 2016; Smith et al., 2013) or sometimes in a more anterior location within the intraparietal sulcus (Gilmore et al., 2019; Gordon et al., 2017; Kong et al., 2019; Shirer et al., 2012). Less frequently, the PMN has been reported to include frontal regions at or near the rostral prefrontal cortex (rPFC; Kim, 2013, 2017; McDermott et al., 2009; Otten & Rugg, 2001) and the anterior cingulate and medial prefrontal cortex (collectively referred to as ‘mPFC’ here). These frontal regions were more commonly reported in task-based analyses (Kim et al., 2013; McDermott et al., 2009; Otten & Rugg 2001) than FC estimates, leading to the proposal that the frontal regions are part of a separate frontoparietal network that is recruited alongside the PMN in certain task contexts (e.g., see Chen et al., 2017). However, Gordon et al. (2017) showed that some individuals do display frontal PMN regions in task-free FC estimates when individual differences are specifically considered. In their analysis, some individuals also showed a further PMN region at or near the ramus marginalis of the cingulate sulcus (RMC). In summary, data support that the PMN may include multiple regions beyond the posteromedial set (i.e., the PCU, rPCC, IPL, rPFC, mPFC, RMC), but emphasize that considerable individual differences may have led to these regions being missed in group-level estimates (Gilmore et al., 2021).

Recently, advances in functional magnetic resonance imaging (fMRI) have allowed the estimation of brain networks reliably within the individual through repeated scanning (Laumann et al., 2015; Poldrack et al., 2015). The individual-level maps capture idiosyncratic details of the networks present in each individual (Braga et al., 2020; Braga & Buckner, 2017; Gordon et al., 2017; Laumann et al., 2015), which has revealed new insights (e.g., Braga et al., 2019, 2020; Braga & Buckner, 2017; DiNicola et al., 2020; DiNicola & Buckner, 2021; Gordon et al., 2023; Newbold et al., 2020; Salvo et al., 2021; Seitzman et al., 2019; Zheng et al., 2021). We showed that the canonical default network comprises at least two parallel distributed networks, DN-A and DN-B, when networks are estimated within individuals, with the two networks exhibiting side-by-side regions in multiple cortical association zones (Braga and Buckner 2017). Of these networks, only DN-A was found to be activated during a task involving thinking about the past or future, indicating that the involvement of the canonical default network in recollection is restricted to the regions encompassed by DN-A (DiNicola et al., 2020). That study did not consider the PMN; however, other individual-level estimates of the PMN have again emphasized idiosyncrasies. Zheng et al. (2021) analyzed extensively collected data from 10 individuals and found that, while all individuals displayed a PMN that included the PCU and rPCC regions, only 4 displayed the mPFC regions, and 5 displayed the RMC region (see ‘Medial Parietal’ network in the first supplemental figure in Zheng et al. 2021; lateral view not shown). In a separate analysis of this same data, Gordon et al. (2017) detected the mPFC PMN regions in 2 individuals, RMC regions in 3 individuals, and rPFC regions in 5 individuals. Notably, in most individuals the cortical patches of mPFC where the PMN regions would have been were ascribed to the salience network (SAL; Dosenbach et al., 2007; Seeley et al., 2007). These prior accounts show that even when more powerful individual-level analyses are deployed, the detection of PMN regions beyond the posteromedial set has been inconsistent.

Although individuals might truly vary in the number of brain regions connected to the PMN, a simpler explanation is that small regions may have been missed in some individuals because they fall below the resolution limits of the imaging procedures. Recently, using a multi-level Bayesian parcellation that incorporates information about individual variability to stabilize network estimation, Kong et al. (2018) defined the PMN as a widely distributed network, including regions in a total of 7 distinct cortical zones, including PCU, rPCC, RMC, rPFC, IPL and mPFC, as well as the anterior insula (aINS; see ‘Control C’ network in Kong et al., 2019). In each zone, the PMN regions consistently bordered the DN, FPN and SAL networks. The aINS regions, while surprising, were also present in 2 individuals in the analysis by Gordon et al. (2017) upon close inspection. Hence, these data support that the PMN is a widely distributed network, extending far beyond the core parietal set, and better matching the expected distributed organization of association networks.

We hypothesized that high-resolution network estimation within individuals would resolve multiple PMN regions beyond the posteromedial set consistently, including in regions that are difficult to resolve such as the insula (Amiez et al., 2016). Our prior work has shown that performing individualized network estimation using high-resolution, high-field 7T fMRI data provides advantages over 3T fMRI, including the separation of distinct networks within the same sulcus, and the detection of smaller regions not observed at lower field strength (Braga et al., 2019). Here, we analyzed data from six extensively sampled individuals who provided high-resolution 7T fMRI data as part of the Natural Scenes Dataset (NSD; Allen et al., 2021). The aim was to identify features of the PMN that were consistent across individuals. We then studied how the PMN relates to other association networks, focusing on nearby networks within the canonical default (DN-A, DN-B; Braga et al., 2017), frontoparietal control (FPN-A, FPN-B; (Braga & Buckner, 2017; Dixon et al., 2018; Thomas Yeo et al., 2011), and salience and cingulo-opercular (SAL; CON; Dosenbach et al., 2007) network regions. To presage the results, we found that in all individuals the PMN displayed regions in upwards of 9 cortical zones, including PCU, rPCC, RMC, IPL, rPFC, mPFC, aINS, as well as potentially dorsal prefrontal cortex (dPFC) and ventromedial prefrontal cortex (vmPFC). A further region may be present in the lateral temporal cortex near to sites suffering from susceptibility artifacts. In each zone, we saw close interdigitation but clear separation of the PMN from adjacent networks. Results were replicated and triplicated in left-out data from the same individuals, and further replicated in an independent dataset comprising eight individuals using high-signal, multi-echo fMRI at 3T. The presence of these regions indicates that the PMN is another instance of a parallel distributed association network, likely formed through a process of specialization and fractionation of an expanded prototypical network motif that is characteristic of association cortex (DiNicola & Buckner, 2021).

## Materials and Methods

### Overview & participants

The Natural Scenes Dataset (NSD; Allen et al., 2021) is a publicly available dataset that comprises high-resolution, high-field 7T fMRI data from eight extensively sampled individuals. Each participant provided approximately 30-40 fMRI sessions, with an average of 2 hours of resting-state (passive fixation) and 38.5 hours of active task fMRI data per participant. MRI sessions were collected approximately once per week. Two participants (S5 and S8) were excluded from our analysis due to excessive head motion during resting state runs, resulting in a final sample of six participants (S1-S4, S6, and S7; age range 23-30 years). Data from each participant were divided into a discovery dataset, for exploratory analysis, and replication and triplication datasets for validation (see Supp. Table S1). The PMN was initially defined in the discovery dataset from each participant using a manually selected seed-based approach, followed by data-driven clustering. We also defined 6 other *a priori* selected networks, including DN-A, DN-B, FPN-A, FPN-B, SAL, and CON, chosen based on their theoretical relevance and spatial proximity to the PMN along the cortex. We then statistically tested the separation of the PMN from these adjacent networks in the left-out, validation datasets. Following statistical testing, we replicated and triplicated the organization of the PMN in the left-out datasets. Finally, to confirm the functional properties of the PMN we assessed the networks for the ‘repetition enhancement’ effect by comparing activity elicited by viewing repeated images in the NSD dataset.

### Resting-state fMRI

Two resting-state fMRI runs were collected per session, one before and one after the main NSD tasks (further details in Allen et al. 2021). Each resting-state run lasted 5 minutes, and a total of 100-180 minutes of resting-state data were acquired over 10-18 sessions for each participant. During the first resting-state run of each session, participants were told to remain awake and fixate their gaze on a centrally presented white crosshair. In the second resting-state run, participants were presented with a red crosshair at the beginning and instructed to take a deep breath when the crosshair turned red. After this cued breathing period, which occurred only once, participants were instructed to fixate. Both types of runs were treated as resting-state data here and counterbalanced first and second runs were allocated to each dataset (discovery, replication, triplication).

### MRI data acquisition, processing and quality control

Functional images were collected using a 7T Siemens Magnetom MR scanner at the Center for Magnetic Resonance Research at the University of Minnesota. Blood-oxygenation-level-dependent (BOLD) images were collected using gradient-echo echo-planar imaging (EPI) at 1.8-mm isotropic resolution with whole-brain coverage with the following parameters: TR = 1,600 ms, TE = 22.0 ms, Flip angle 62 degrees, FOV = 216 mm (FE) × 216 mm (PE), slice thickness 1.8 mm, slice gap 0 mm, matrix size 120 × 120, echo spacing 0.66 ms, bandwidth 1,736 Hz per pixel, partial Fourier 7/8, iPAT 2, multiband slice acceleration factor = 3, and 84 slices acquired in the axial plane. Dual-echo fieldmaps were collected for post hoc correction of EPI spatial distortion. Pre-processed versions of the data are shared in the NSD (http://naturalscenesdataset.org), which include steps to correct for slice acquisition timing, alignment of data from each TR to correct for head motion within a run, alignment across sessions, correction for EPI distortion, all performed within one interpolation step. Detailed information on preprocessing procedures can be found in Allen et al., 2021).

We performed quality control on the NSD resting-state data and excluded runs that did not pass rigorous criteria for head motion. Whole runs were automatically excluded if maximum framewise displacement (FD) was greater than 0.4 mm, or maximum absolute motion was greater than 2.0 mm. We also visually inspected any runs with maximum FD > 0.2 mm or a maximum absolute motion > 1 mm, and excluded any that exhibited visible movement. This resulted in a total of between 6-35 resting-state runs per participant (S1: 35 runs; S2: 6; S3: 16; S4: 12; S6: 19; and S7: 18). For the five participants with 12 or more included runs, the data were divided into two or three groups: a discovery dataset plus replication and triplication datasets (see Supp. Table S1).

For functional connectivity analysis, we performed additional preprocessing on the resting-state data following procedures outlined in Braga et al. 2019. Nuisance variables were regressed out, including six parameters to account for head motion, as well as whole-brain, ventricular, and deep white matter signal, and temporal derivatives. Nuisance regression was performed using 3dTproject (AFNI version 2016.09.04.1341; Cox 1996) on native-space-projected BOLD data resampled to 1mm isotropic resolution (i.e., the ‘func1pt0mm’ version of the NSD data). Data were bandpass filtered at 0.01–0.1Hz (using 3dBandpass from AFNI). Next, we projected the data onto a standardized cortical surface containing 163,842 vertices (fsaverage7) per hemisphere using FreeSurfer’s vol2surf command (Fischl et al., 1999a) and smoothed along the surface using a 2.5mm FWHM kernel. The highest resolution cortical mesh was used to minimize blurring and preserve fine-scale distinctions between networks. The smoothing kernel was chosen by eye based on preliminary analyses on one individual, carefully assessing the trade-off between minimizing smoothing (i.e., preserving details), maximizing correlation values, and minimizing noise or ‘speckling’ in randomly chosen seed-based correlation maps in the exploratory data. Functional connectivity matrices were estimated in each participant by computing vertex-vertex Pearson’s product-moment correlation for each run, z normalizing, averaging across runs within each dataset in each individual, then converting back to r values. These matrices were then used for network estimation (Braga & Buckner 2017) for manual seed-selection using the Connectome Workbench (Marcus et al., 2011) and for data-driven clustering.

### Seed-based functional connectivity analysis

Our initial analyses sought to identify the PMN within each individual, anchoring on the spatial distribution of key component regions previously reported (Gilmore et al., 2015; Shirer et al., 2012). The two key components were the prominent PCU and rPCC regions that most consistently comprise the PMN (Gilmore et al. 2021). We initially selected seeds in the lateral PFC. This was done (i) to align with our previous seed-based analyses targeting other association networks within individuals (Braga & Buckner 2017; Braga et al., 2020; DiNicola et al., 2020), (ii) to allow long-distance correlation patterns to be appreciated (e.g., at our key component regions) without the confound of local blurring near the seed, and (iii) following initial observations that the PMN could be reliably defined from seeds in the PFC. The organization of the PMN was then confirmed by targeting seeds to five cortical zones in each individual. The zones include the rPFC, aINS, IPL, posteromedial cortex (posterior cingulate, precuneus, cuneus and retrosplenial cortices), and mPFC (Fig. 1). These seeds replicated the detailed organization of the PMN (correlation maps thresholded at r > 0.2), including confirmation of a PMN region in the rPFC (Fig. 2). We also targeted six other networks, DN-A, DN-B, FPN-A, FPN-B, SAL, and CON, with seeds selected in the same 5 cortical zones (with the exception of DN-A, for which no region could be found in the aINS zone; Supp. Fig. S3). These seeds were used to statistically test for a dissociation in the correlation between the networks in each individual using the left-out datasets.

**Figure 1:**
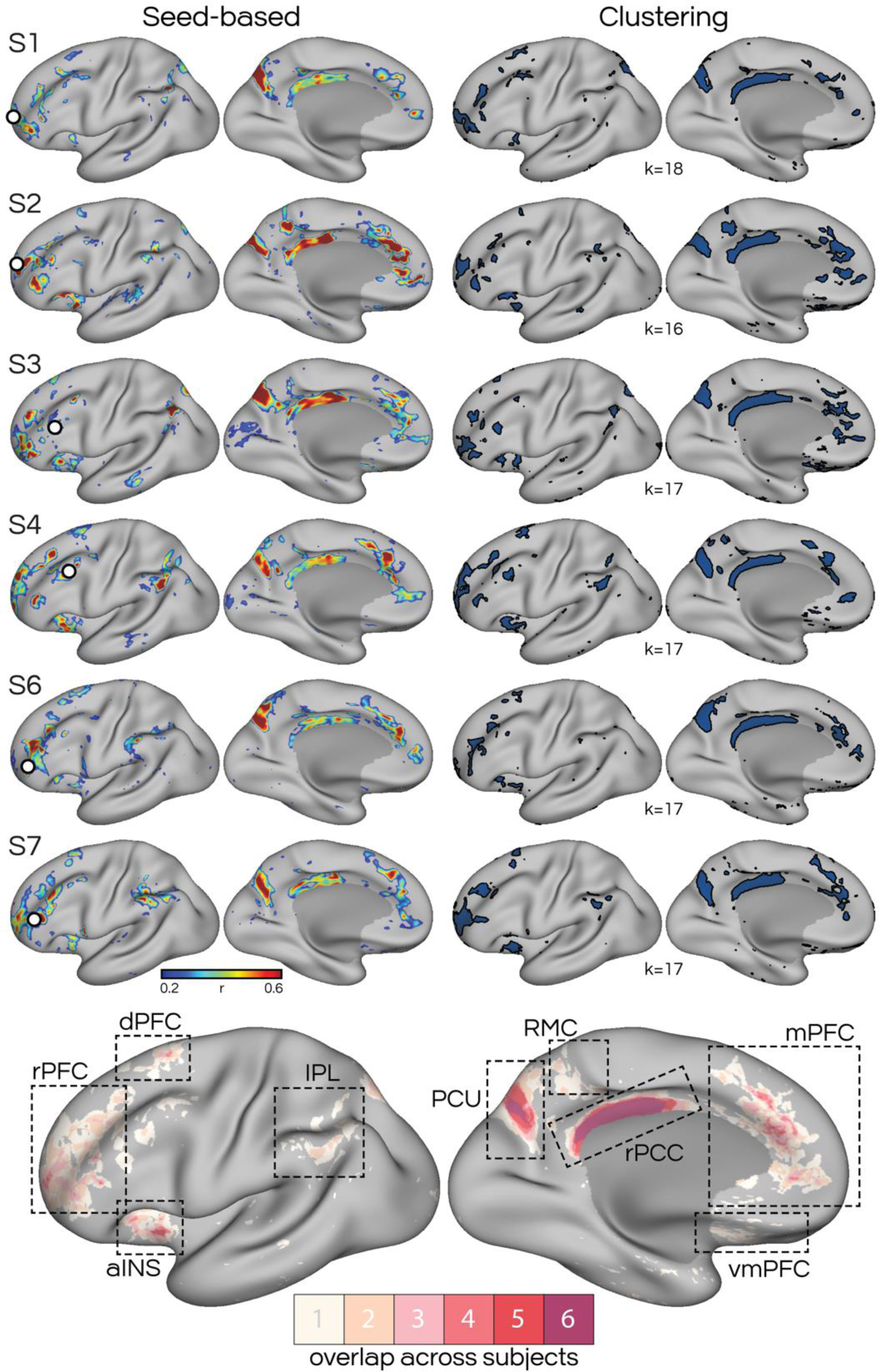
High-field, high-resolution fMRI functional connectivity reveals a parietal memory network (PMN) that includes smaller regions distributed across multiple cortical zones. The PMN was estimated using seed-based and clustering approaches. Top panel) The left column shows lateral and medial views of seed-based functional connectivity estimates of the PMN for all six individuals. Seeds were selected from the prefrontal cortex in each individual (S1-S7, rows) and marked with white circles. The right column represents the PMN estimated using a clustering approach (Kong et al. 2019). Lower panel) An overlap map of the PMN was calculated for all six individuals, using the estimate of the PMN from the clustering analysis (right column, top panel). Regions are color-coded according to the number of individuals that displayed a region at each vertex. The precuneus (PCU) and rostral posterior cingulate cortex (rPCC, see dashed boxes) regions of the PMN showed extensive overlap across all 6 individuals, whereas other regions showed more variability. rPFC; rostral prefrontal cortex, dPFC; dorsal prefrontal cortex, aINS; anterior insula, IPL; inferior parietal lobule, RMC; ramus marginalis of the cingulate sulcus, mPFC; medial prefrontal cortex, vmPFC; ventromedial prefrontal cortex.

**Figure 2:**
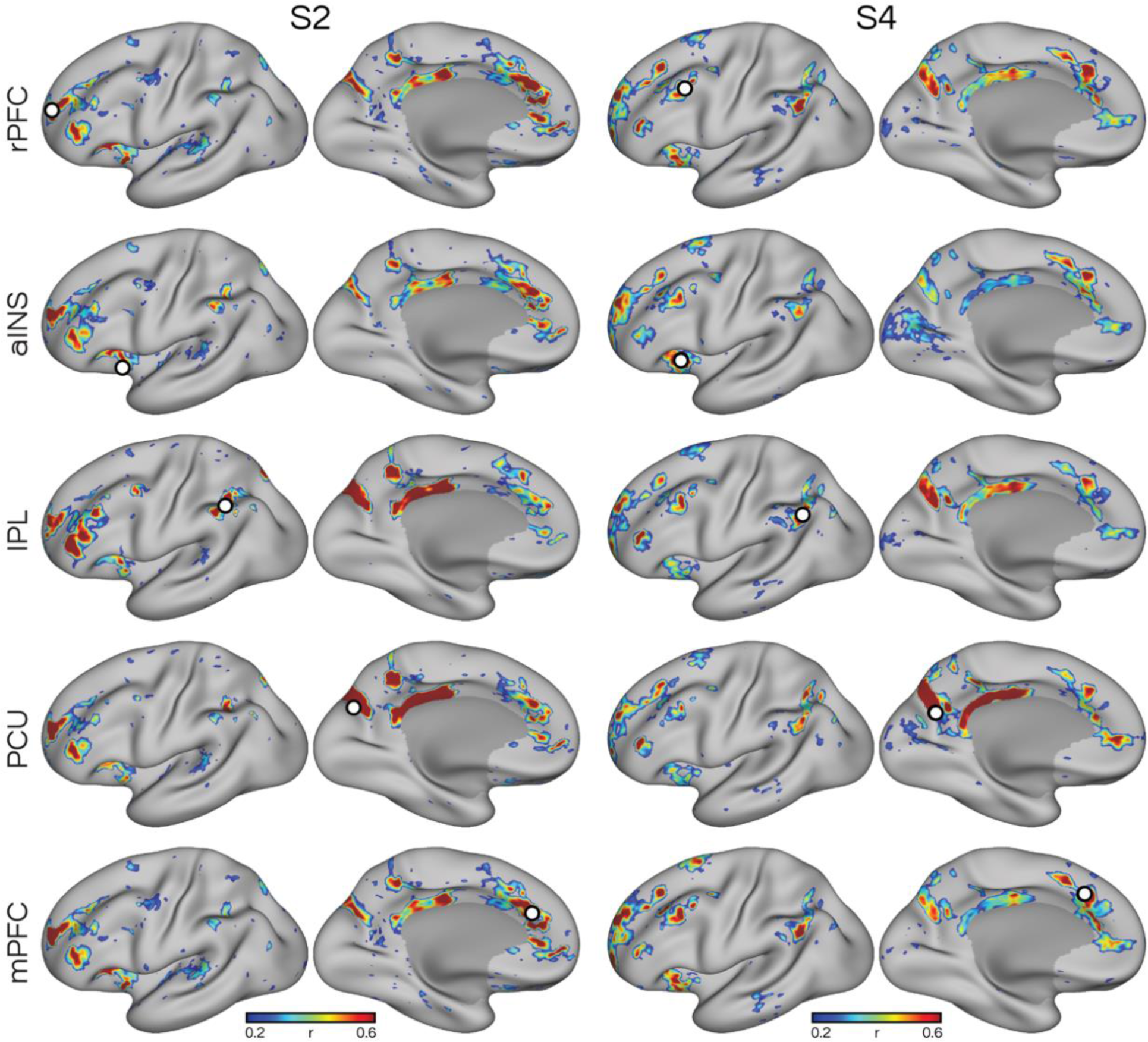
Seed-based functional connectivity at high resolution reproducibly defines the parietal memory network (PMN) within individuals. Five estimates of the PMN are shown in two representative participants (S2 and S4). Seeds (white circles) were selected from five cortical zones (rows), targeting converging estimates of the PMN. In each case, the same distributed network was observed, extending beyond the core set of posterior midline regions in the precuneus (PCU) and rostral posterior cingulate cortex (rPCC). The PMN consistently included regions in the rostral prefrontal cortex (rPFC), anterior insula (aINS), inferior parietal lobule (IPL), and medial prefrontal cortex (mPFC), and could be defined from seeds placed in each of these zones. The converging estimates reinforce that the PMN includes regions distributed throughout multiple cortical zones.

### Clustering approach

A multi-session hierarchical Bayesian model (MS-HBM) parcellation method (Kong et al., 2021) was employed for confirmation of network organization. This approach provides individual-specific network estimates by integrating priors from multiple levels (e.g., group atlas, cross-individual and cross-run variation) to stabilize network estimates. The MS-HBM parcellation method was applied to define networks using the discovery dataset. In each individual, we used a *k* value (i.e., number of clusters) between 14–20 and selected the solution that best matched the networks observed in the seed-based analysis, respecting that the same value of *k* can over-split or over-lump networks in different individuals. Namely, the lowest value of *k* that separated the PMN from other networks – with an initial focus on PMN’s separation from FPN-A and FPN-B – as observed in the seed-based approach was taken as the solution. Notably, to achieve separation of the PMN and match our seed-based observations, in some subjects a level of *k* was used that over-split our expected breakdown of DN-A (i.e., note diminished DN-A region in the retrosplenial cortex in subjects S1, S6 & S7 in Supp. Fig. S6). The highlighted details in Supp. Fig. S6 allow appreciation of the differences between seed-based and clustering solutions of DN-A in these subjects. Note that in all subjects the clustering estimate of the PMN overlapped closely with seed-based estimates (see multiple details in Figs. 4-5 and Supp. Figs. S6-S9), indicating that this over-splitting of DN-A did not affect the estimate of the PMN. The clustering analysis provided a data-driven confirmation of the patterns observed in the manual seed-based approach, while minimizing observer bias. The two approaches provided converging solutions and confirmed the PMN as a distributed network that is distinct from surrounding networks (Fig. 1).

### Statistical analysis

Seeds targeting each network in each cortical zone and individual were selected using the discovery dataset and used to extract timeseries for each run in replication datasets in subjects that provided enough data (see *MRI data acquisition, processing and quality control*). Pearson’s product-moment correlations were calculated between the extracted timeseries for each run in the replication dataset. We calculated correlations across all 34 seeds (5 seeds in each network except DN-A, which has only 4 seeds), resulting in a 34 by 34 FC matrix for each run and each individual. The elements in these seed-wise FC matrices were then averaged together to generate network-network correlation values for each run. For within-network FC, we averaged only the lower triangle of the symmetric matrix to avoid repeats. A paired t-test was performed to compare within-versus between-network FC (Fig. 7 and Supp. Fig. S12). Six separate t-tests (e.g., comparison between PMN vs. DN-A, PMN vs. DN-B, PMN vs. FPN-A, etc) were conducted for each target network. Benjamini-Hochberg correction was used for the six comparisons. Following these individual-level analyses, we tested for robustness at the group level by averaging network-wise FC across sessions for each individual and then comparing within-versus between-network FC, as in the individual-level analyses.

### Continuous recognition task

We sought to confirm that the PMN identified here displayed functional properties characteristically ascribed to the PMN (e.g., Gilmore et al., 2015). Namely, the PMN shows a repetition enhancement effect, where the perceived familiarity of a stimulus (e.g., an image) is associated with increased responses. This response includes a ‘flip’ from below-baseline activity during initial presentation of novel images, to above-baseline increased activity for repeated presentations. To confirm the functional characteristics of the PMN, task data from the NSD experiment, a continuous recognition task, were analyzed. Participants viewed a series of color natural scene images and were asked to respond every time they saw an image while undergoing scanning. Participants were instructed to press a button with their right index finger if they thought the presented image was new or press another button with their right middle finger if the presented image had been shown previously. Each run included 62-63 trials, with an image presented every 3 seconds, followed by 1s fixation period. Twelve runs were collected in each session, yielding a total of 750 trials per session. Each participant underwent 30-40 sessions over one year (S1: 40 sessions; S2: 40; S3: 32; S4: 30; S6: 32; and S7: 40; though the last 3 sessions from each participant had not yet been released and were not analyzed here). More details on the task design are provided in Allen et al., 2021. The large number of trials available (ranging from 22,500 to 30,000 trials per participant) allows reliable exploration of repetition effects. The experiment consisted of 10,000 distinct images, each of which was presented up to three times throughout the experiment, depending on the number of completed sessions by a participant. Participants also completed a variety of behavioral measures, a final memory test, and an image-similarity assessment after the scan, not analyzed in the present study.

The NSD public release includes beta maps for each trial of the continuous recognition task (representing the percent BOLD signal change evoked by each trial relative to a baseline), as well as mean FD and voxel-wise tSNR for quality control purposes. We excluded runs with mean FD greater than 0.16 mm and tSNR lower than 20. We compared betas (beta version 3 provided in the NSD) from trials containing repeated versus novel presentations of the same images, restricting the analysis to correct responses only. To focus on short-term recognition memory, and avoid the increased variance of comparing data across sessions, we considered only images that were presented 3 times within the same session. The within-session repeats were also more likely to be recognized as familiar (within-session hit rate = .98, across-session hit rate = .72, p < .001). By design, participants were presented with the ‘new’ condition more frequently than the ‘old’ condition during the initial sessions, and were presented with the ‘old’ condition more often during the later sessions. To avoid odd-ball effects, we only included sessions where the difference between the ‘new’ and ‘old’ conditions had a ratio less than 0.3. In other words, only sessions in which both trial types were relatively balanced, with neither trial type comprising less than 35% or more than 65% of the total trials, were included in the analysis. As a result, 14 sessions were included in the analysis for each individual. Similar results were obtained in an initial analysis that included all sessions.

Trials were divided into three types relating to first (P1), second (P2), and third (P3) presentation. Only trials which were correctly identified as novel in their first appearance (i.e., correct rejections) and correctly identified as repeats in the second and third appearances (i.e., hits) were included in the analysis. We performed two-tailed t-tests comparing each pair of trial types to obtain a statistical map for each comparison of trial types. To control for the potential confounding effects of response time (RT), we included RT as a covariate (Taylor et al., 2014; Yarkoni et al., 2009). Results did not differ considerably when RT was not modelled. All statistical analyses were conducted using FSL’s Permutation Analysis of the Linear Model (PALM; Winkler et al., 2016) in MATLAB 2018b. A permutation-based correction for multiple comparisons was performed across all vertices to control the family-wise error rate with p < 0.05 (1,000 permutations). To specifically test whether the PMN and other networks showed the repetition enhancement effect, we took the manually selected seeds targeted to the PMN, DN-A and DN-B for the resting-state statistical analysis, calculated an average beta from these seeds per network, and performed a t-test (two-tailed). We also computed mean beta values within network regions by averaging vertex-wise betas within network regions estimated using the clustering analysis, and performed a t-test between conditions. We repeated the same procedure for each network. Bonferroni correction for multiple comparisons was applied for the three pairs of trial types (critical value = 0.05/3).

### Validation in an independent dataset

#### Overview & participants

We validated the observed organization of the PMN in an independent 3T MRI dataset containing 8 extensively sampled participants collected at Northwestern University. All individuals were native English speakers, neurologically healthy, and had normal or corrected-to-normal vision (4 females, age range 22-36 years, mean age = 26.8 ± 5.3 years). Participants provided written informed consent in compliance with procedures approved by the Northwestern University Institutional Review Board and were paid for their participation. The experiment consisted of eight sessions. During each session, participants completed two 7-min resting-state (passive fixation) runs which were collected as the first and final runs in each session. This resulted in a total of 112 minutes (2 runs x 8 sessions x 7 min) of resting-state data per participant. Participants completed six to eight active tasks between the two resting-state runs in each session, but we analyzed only the resting-state data in the present study.

Data were quality controlled and runs that did not pass the same criteria as the NSD data for head motion were excluded. This led to a total of between 10-16 resting-state runs per participant (S1: 16 runs; S2: 16; S3: 16; S4: 16; S5: 10; S6: 15; S7: 14; and S8: 14). For the seven participants with more than 10 good quality runs, the data were divided into a discovery dataset and a replication dataset. Only the discovery dataset was used for the present analyses (see Supp. Table S2).

#### MRI data acquisition, processing and quality control

MRI data were collected at the Center for Translational Imaging at Northwestern University on a 3T Siemens Prisma scanner. A high-resolution T1-weighted magnetization-prepared rapid acquisition gradient echo (TR = 2,100 ms, TE = 2.9 ms, FOV = 256 mm, flip angle = 8°, slice thickness = 1 mm, 176 sagittal slices parallel to the AC-PC line) was acquired after the first resting-state run. Functional MRI were collected using a 64-channel head coil with a multi-band, multi-echo sequence with the following parameters: TR = 1,355 ms, TE = 12.80 ms, 32.39 ms, 51.98 ms, 71.57 ms, and 91.16 ms, flip angle = 64°, voxel size = 2.4 mm, FOV = 216 mm x 216 mm, slice thickness = 2.4 mm, multiband slice acceleration factor = 6. Functional MRI data were pre-processed using the iProc pipeline (Braga et al., 2019) with the following steps. Runs with excessive head motion (a maximum FD > 0.2 mm or a maximum absolute displacement > 1 mm) were visually inspected and excluded if they exhibited noticeable movement. The first nine volumes (approximately 12 seconds) were removed to allow for T1 attenuation, and a mean BOLD template was generated using the remaining runs. Brain extraction was performed using FSL’s Brain Extraction Tool (FSL v6.0.3). Nuisance signals relating to deep white matter, ventricular, and whole brain signal time series were regressed out of the data, followed by bandpass filtering at 0.01-0.10 Hz. Data were then projected onto a high-resolution standard surface mesh (fsaverage6, 40,962 vertices per hemisphere) using Freesurfer (Fischl et al., 1999). Finally, the data were spatially smoothed with a 2.5 mm full width-half maximum smoothing width, optimized to maintain precision while excluding noise. Pearson’s product moment correlations were computed pairwise between vertices to generate a correlation matrix.

#### Clustering approach

The same MS-HBM parcellation method was applied to extract clusters (Kong et al., 2021) in the discovery dataset. In each individual, a *k* value between 14–20 was used, and the solution that best matched the networks observed in the seed-based analysis was selected.

## Results

### High-resolution fMRI reveals that the PMN is distributed across multiple cortical zones

We quantified data quality by calculating temporal signal-to-noise ratio (tSNR) for each participant, which revealed high average tSNR in each individual (Supp. Table S1), despite the small voxel size (1.8 mm isotropic; see Supp. Fig. S1). Runs passing quality control were divided into discovery and validation (replication and triplication) datasets in each individual. We initially explored the discovery dataset, and then performed hypothesis-testing analyses in the validation datasets, before replicating and triplicating network maps in the validation datasets.

On initial exploration of functional connectivity patterns in one of the NSD subjects, the observation was made that a network resembling the PMN could be defined by selecting seed regions from the posteromedial cortex. The estimated network included prominent regions in characteristic PMN locations, including the PCU and rPCC (Gilmore et al., 2015; Power et al., 2011; Thomas Yeo et al., 2011), but also included regions in multiple lateral and frontal cortical zones, including rPFC, dPFC, IPL, RMC, mPFC, and aINS (Fig. 1). The resulting network appeared as a distributed network with regions in multiple cortical association zones. Motivated by this, we explored the organization of this network in other individuals, using multiple converging approaches. First, we manually selected seeds from the rPFC in each individual (Fig. 1, left column). In each participant, the PMN appeared as a distributed network with regions in multiple cortical zones. Next, we confirmed the stability of the key regions by defining the PMN using seeds in five cortical zones (Fig. 2). These analyses recapitulated the PMN as a distributed network that could be defined from seeds in multiple cortical zones, including regions underemphasized in the literature such as the aINS and mPFC (Fig. 2). We next confirmed that we were able to define other networks in the proximity of the PM – including DN-A, DN-B, FPN-A, FPN-B, SAL, and CON – to ensure that the estimated PMN was not being confused with one of these networks. In each participant, we were able to define all other networks using seeds in the lateral prefrontal cortex, and to confirm that the PMN was a distinct network (Supp. Figs. S3–S5). Once all networks had been defined using a manual seed-based approach. we performed a data-driven clustering (Kong et al., 2019) to define all the networks simultaneously while reducing observed bias (Fig. 1, right column, and Fig. 3). This analysis confirmed, using a different analytic approach, that when defined in individuals using high-resolution data the PMN includes regions in upwards of 9 cortical zones (Fig. 1).

**Figure 3:**
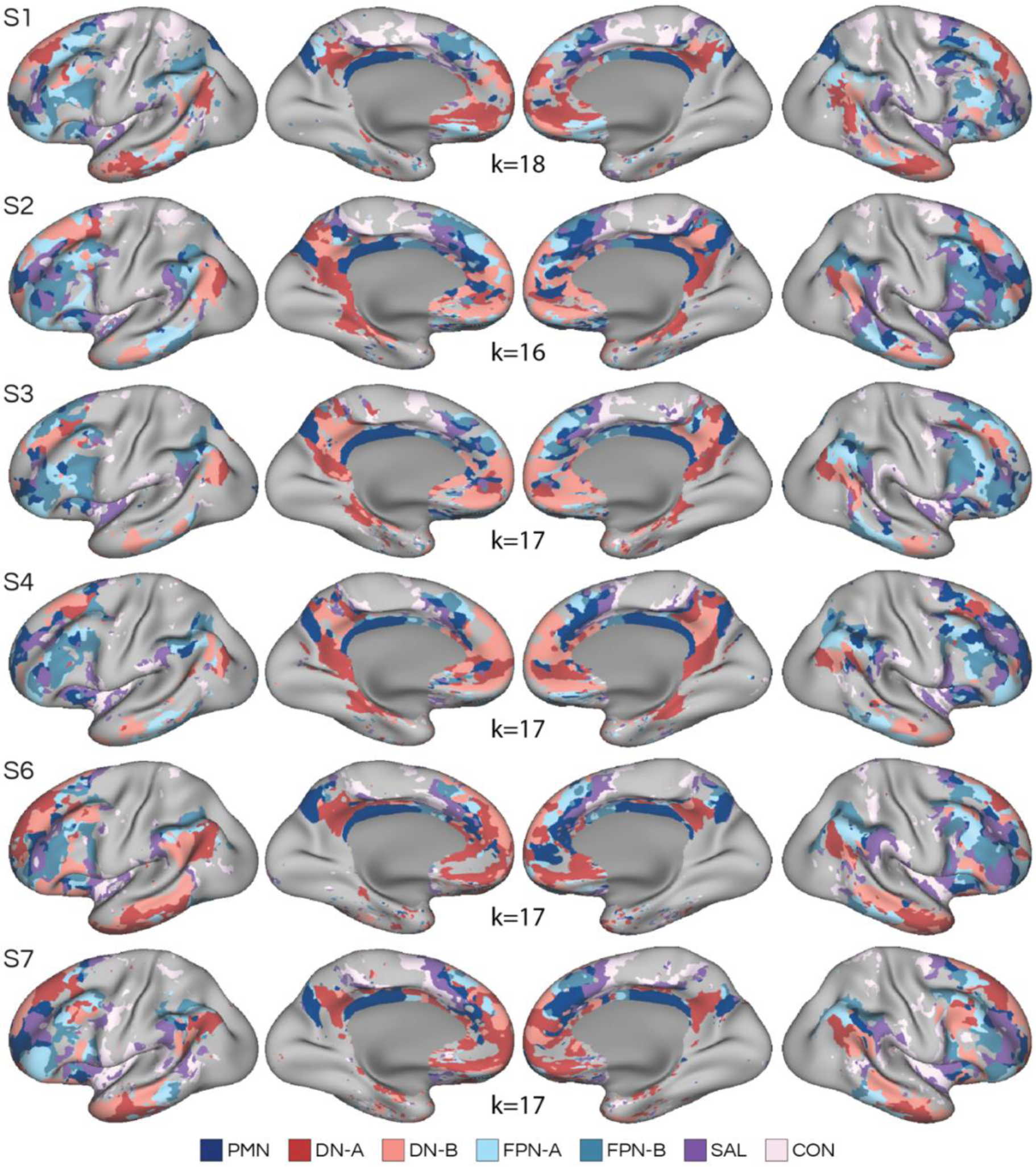
Data-driven clustering situates the parietal memory network (PMN) alongside multiple parallel distributed networks. Functional connectivity was used to estimate the organization of 7 a priori selected networks using a multi-session hierarchical Bayesian model (Kong et al., 2019). Networks were identified from the clustering solution based on anatomical distinctions reported in the literature (Braga and Buckner 2017; Braga et al. 2019; 2020; Dosenbach et al., 2007; Seeley et al., 2007). In multiple cortical zones, the PMN was located adjacent to the same set of networks, emphasizing a fine-scale interdigitated organization of parallel distributed networks. The *k* value used ranged from 16-18 networks, based on the lowest value that separated the full set of a priori selected networks in each participant. Networks included default network A (DN-A), default network-B (DN-B), frontoparietal control network A (FPN-A), frontoparietal control network B (FPN-B), salience network (SAL) and cingulo-opercular networks (CON).

A strong possibility, as suggested by other authors (Gilmore et al., 2021; Gordon et al., 2017), is that individual differences in network anatomy may have obscured detection of these multiple PMN regions, particularly in prior analyses that relied on group averaging. To explore this, we computed an overlap map for our 6 individuals by taking the binary estimate of the PMN from the clustering analysis (Fig. 1, right column), and calculating how many subjects displayed a PMN region at each surface vertex in a common (i.e., fsaverage7) space (Fig. 1, bottom). This analysis revealed that the prominent PCU and rPCC regions of the PMN showed extensive overlap across all 6 individuals, whereas other regions were more variable. Notably, the aINS region also showed high overlap across most subjects. It is possible that this aINS region may have been missed in prior analyses due to the difficulty of resolving insular anatomy, underscoring the need for high-resolution approaches (e.g., Amiez et al., 2016). The mPFC region showed a moderate degree of overlap across individuals, while other regions were less consistent.

As a final confirmation, after all statistical analyses had been performed we replicated and triplicated the definition of the PMN in left-out runs for individuals with sufficient data (Supp. Fig. S14). These replications provide further evidence that the PMN reliably contains multiple regions beyond the core posteromedial set.

### The PMN is distinct from networks within canonical default, frontoparietal control and salience/cingulo-opercular regions

Conflicting accounts of the functional role and anatomical organization of the PMN have been proposed, which could be clarified by considering how the present definition of the PMN relates to other networks. The PMN has been linked to canonical default and frontoparietal control networks through analysis of resting-state functional connectivity (e.g., Barnett et al., 2021; Cooper et al., 2021; Cooper & Ritchey, 2019; Doucet et al., 2011; Power et al., 2011; Thomas Yeo et al., 2011) and the mPFC regions we observed here are typically ascribed to the salience and cingulo-opercular networks (Dosenbach et al., 2007; Seeley et al., 2007; though see Kong et al. 2019). To understand how the PMN relates to these systems, we zoomed in on specific cortical zones where the networks have previously been conflated and compared the location of PMN to other network regions (Figs. 4-5 & Supp. Figs. S6-S9).

**Figure 4:**
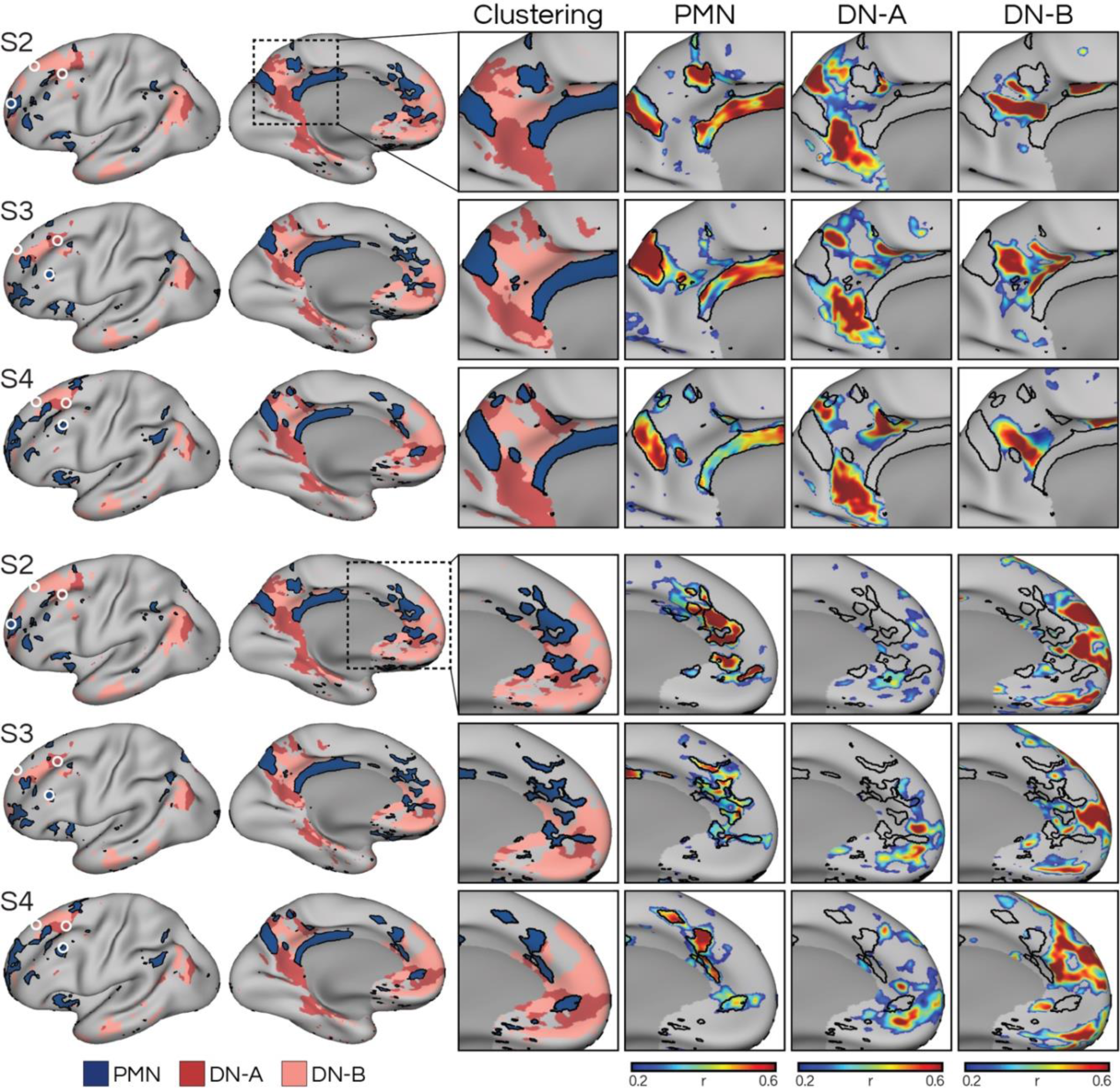
Detailed anatomy of the parietal memory network (PMN) reveals regions that are closely-knit but distinct from networks within the canonical default network (DN). Left column shows the clustering-defined networks from Fig. 3 and the location of manually selected seeds (white circles) initially used to define the networks (see Supp. Fig. 3). Right panels show a zoom-in of the posteromedial (top three rows) and medial prefrontal (bottom rows) cortex, showcasing the detailed interdigitation of network regions. The PMN occupied distinct regions that were closely juxtaposed with regions of default network A (DN-A) and B (DN-B). The black lines represent the boundaries of the PMN calculated from the clustering approach, to serve as landmarks for comparing across panels. Three representative participants (S2, S3, and S4) are shown, with the remaining 3 shown in Supp. Fig. S6.

**Figure 5:**
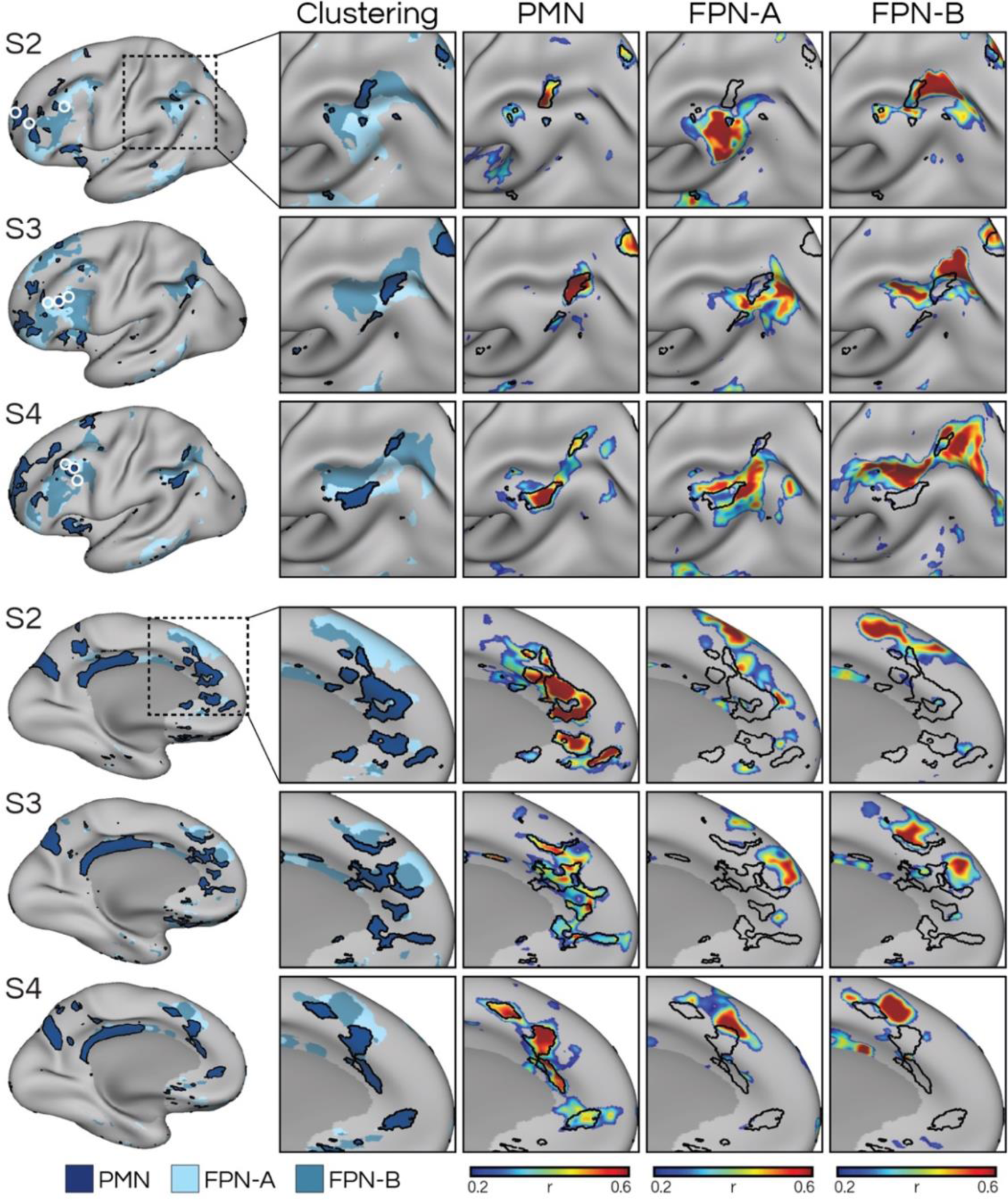
Detailed anatomy of the parietal memory network (PMN) reveals regions that are closely-knit but distinct from networks within the canonical frontoparietal control network (FPN). Left column shows the clustering-defined networks from Fig. 3 and the location of manually selected seeds (white circles) initially used to define the networks (see Supp. Fig. S4). Insets show a zoom-in of the intraparietal sulcus (top three rows) and medial prefrontal cortex (bottom rows), showcasing the interdigitation of network regions. The PMN occupied distinct parts of the cortex that were closely juxtaposed with regions of frontoparietal control network A (FPN-A) and B (FPN-B). Three representative participants (S2, S3, and S4) are shown, with the remaining 3 shown in Supp. Fig. S7.

The exact location and shape of the PMN regions varied appreciably across individuals, but broad consistencies could be observed. In Figs. 4-5 and Supp. Figs. S6–S9, we zoom in on selected cortical zones to show how the PMN is juxtaposed with but is distinct from other networks, displaying a complex fine-scale organization. In the posteromedial cortex (Fig. 4 and Supp. Fig. S6, top panel), the PMN includes multiple regions that surround but are strikingly separable from the regions of DN-A and DN-B (see black outlines of PMN in Fig. 4 insets). In the posterior and middle cingulate cortex, the PMN was consistently located posterior to FPN-A and FPN-B regions (Fig. 5 and Supp. Fig. S7). SAL and CON regions were typically positioned within and/or across the marginal sulcus in the paracentral lobule (Supp. Figs. S8 & S9). Similar juxtapositions were also observed in the medial prefrontal cortex: the regions of DN-A and DN-B were generally positioned in more rostral and ventral sites (Fig. 4 and Supp. Fig. S6), regions of FPN-A and FPN-B were generally in more dorsal locations (Fig. 5 and Supp. Fig. S7), and the regions of SAL and CON were largely in more posterior sites to the PMN (Supp. Figs. S8 & S9). Hence the PMN appears to sit at the confluence of DN-A, DN-B, FPN-A, FPN-B, and SAL in the medial prefrontal cortex, with the CON typically being separated from the PMN by SAL regions. The clustering solution also revealed a similar juxtaposition between the networks in the anterior insular (Supp. Fig. S10); though notably the separation between these networks in the insula was more difficult to achieve in the seed-based analysis (not shown).

A closer correspondence with FPN-A and FPN-B was also seen in the lateral parietal cortex, where PMN regions were sometimes located exactly in between FPN-A and FPN-B regions at or near to the intraparietal sulcus (see zoom-ins in Fig. 5 and Supp. Fig. S7). The intraparietal PMN regions were generally more anterior to previously reported parietal locations in the “posterior inferior parietal lobe” or “dorsal angular gyrus” reviewed in Gilmore et al. (2015; 2021), sometimes extending into supramarginal gyrus and corroborating other accounts (Gordon et al., 2017; Kong et al., 2019; Shirer et al., 2012). The regions belonging to SAL and CON were generally more ventral and anterior to the intraparietal PMN regions (Supp. Figs. S8 & S9). This distinction is notable because the anterior inferior parietal lobe, proximal to the postcentral sulcus, is a characteristic region of CON (Gordon et al., 2017; and see Yeo et al., 2011). Here the PMN is distinguished from both SAL and CON (e.g., see black PMN boundaries in Supp. Figs. S8 & S9).

A further point regards whether the estimated SAL and CON networks were correctly identified, or whether the CON network here instead represents a premotor network (PreM) that surrounds the somatomotor strip (see exploration of this region and relationship to CON in Gordon et al. 2023). For instance, in the solution by Gordon et al. (2017), the salience network occupies mPFC regions that here we ascribe to the PMN. Here, we based our naming on the parcellation of Kong et al. (2019), where “Control C”, “Salience/VenAttn B”, and “Salience/VenAttn A” relate to our PMN, SAL and CON networks here, respectively. To determine whether we had mislabeled the networks, we conducted a targeted analysis to determine whether the PreM network could be defined as separable from CON, and further confirm the separation between all 4 networks using seeds in the anterior midline. Seeds were manually selected from the dorsomedial prefrontal cortex in the three individuals that provided particularly good separation between networks (S2, S3, and S7). In each case, the PMN, SAL, CON, and PreM networks could be defined by the dorsomedial prefrontal seeds, with each network occupying distinct portions of the cortex throughout the brain (Fig. 6). Thus, regardless of the naming convention, our analyses support a separation of three distributed networks, SAL, CON, and PMN, when high-resolution, high-field, 7T data is analyzed. The close juxtaposition between regions of SAL and PMN suggest that these two networks may easily be blurred together in lower-resolution or group-averaged data.

**Figure 6:**
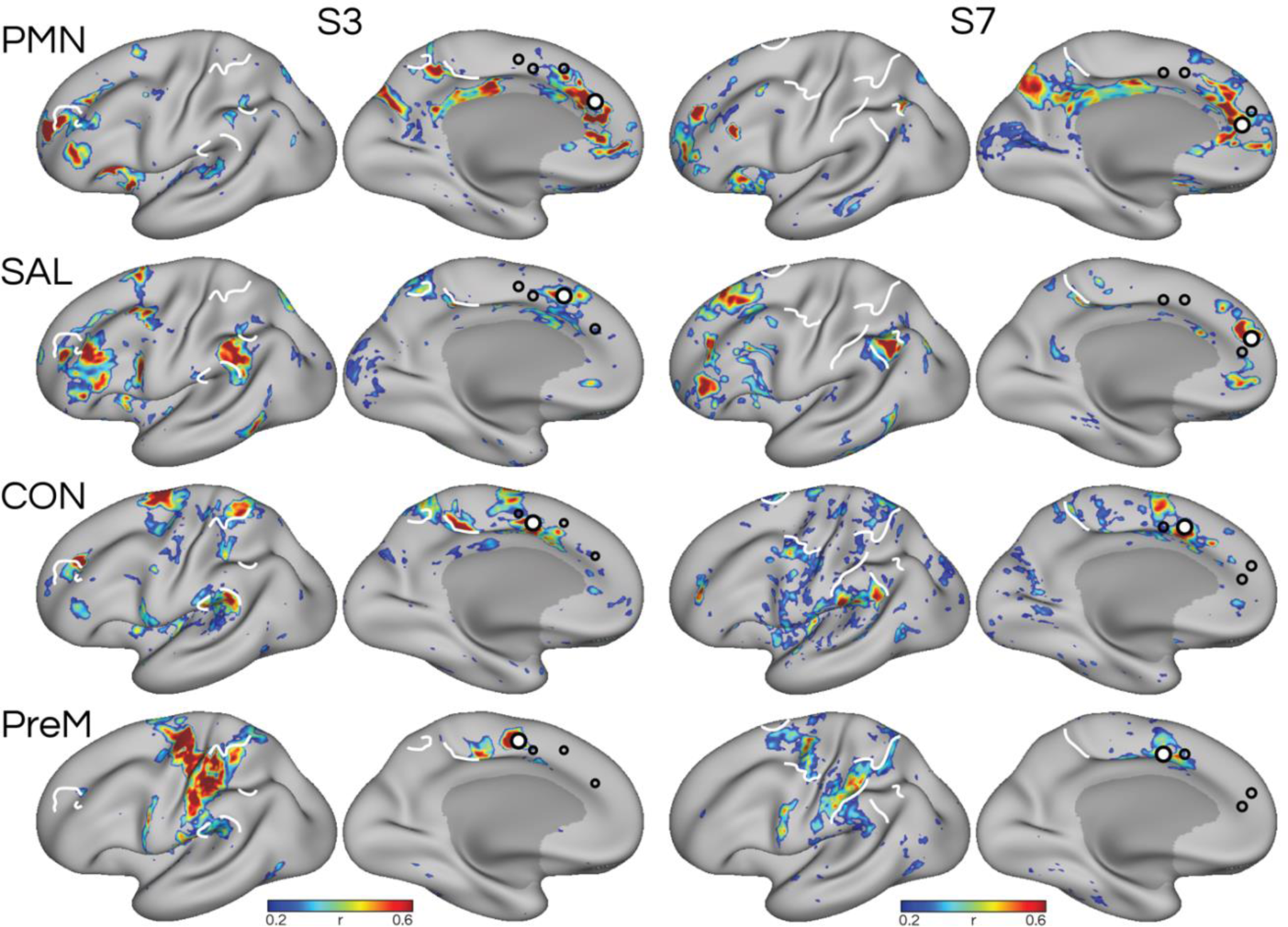
Targeted analysis in two individuals confirms that the parietal memory network (PMN), salience network (SAL), and cingulo-opercular network (CON) are distinct, and are distinct from a premotor network (PreM) that surrounds primary somatomotor regions. Seeds were chosen from the dorsomedial prefrontal cortex in an anterior-posterior progression to target PMN, SAL, CON, and PreM in two subjects (S3 and S7) that provided particularly good separation between the networks in the high-resolution dataset. One additional subject (S2) is shown in Supp. Fig. S18. This analysis showed that SAL and CON are distinguishable from each other, and the premotor network, with each network occupying distinct regions of the cortex. White lines serve as hand-drawn landmarks for comparing across panels. White circles indicate the seed used to define the network shown in that panel, and black hollow circles represent seeds for the other networks shown.

### PMN is statistically and reproducibly dissociated from nearby networks

The observed distinction between the PMN and surrounding networks was statistically tested in the left-out validation datasets. We selected seeds to target each of the 7 *a priori* selected networks (PMN, DN-A, DN-B, FPN-A, FPN-B, SAL, CON) in the posterior and anterior midline, the posterior and anterior lateral surface, and the anterior insula. Within each of these 5 broad regions we were able to select seeds that delineated the all the networks with the exception of DN-A, which did not show a large enough region within the insula for targeting (seed locations shown in Supp. Figs. S3–S5). Seeds were selected using the discovery dataset and tested for dissociation in the validation datasets. Pairwise seed-seed correlations were grouped into within-network and between-network correlations. For all networks tested, within-network correlations were significantly higher than between-network correlations in each individual (Supp. Figs. S11 & S13; all p < .05, Benjamini-Hochberg corrected for 6 networks compared), with few exceptions (see red bars in Supp. Figs. S11 & S13). Importantly, the PMN was statistically dissociated from the other networks (all p < .05, corrected) in all individuals and in all datasets. This result was also significant in a post-hoc group-wise analysis (Fig. 7 and Supp. Fig. S12). These results therefore show that the PMN is statistically dissociable from the other networks including DN-A, DN-B, FPN-A, FPN-B, SAL, and CON using seeds placed in multiple cortical locations. We repeated this analysis in the left-out triplication dataset for the 3 subjects that provided enough data, which confirmed the statistically significant separation between the networks (Supp. Figs. S12 & S13).

**Figure 7:**
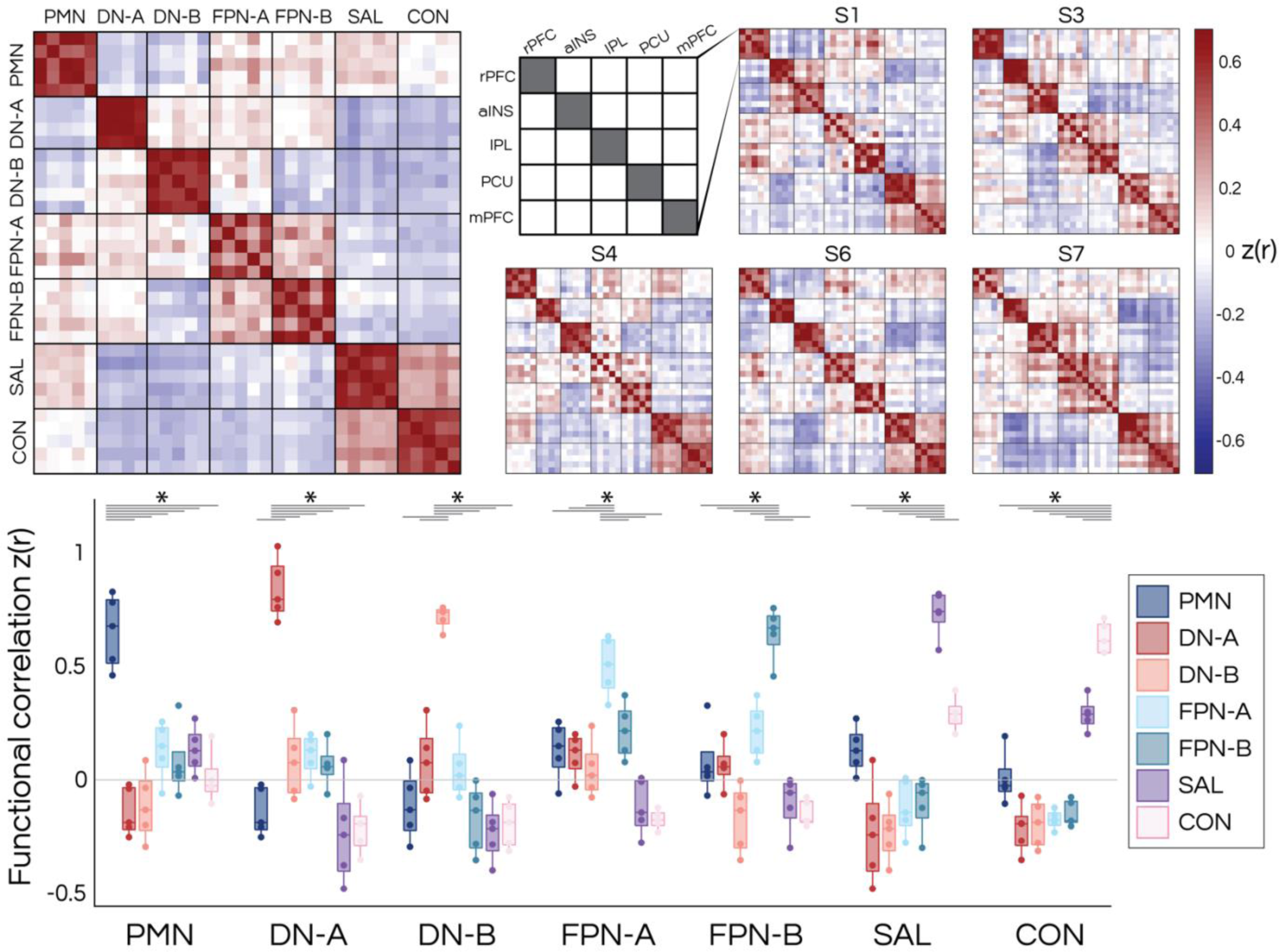
The parietal memory network (PMN) is statistically distinct from nearby networks. In left-out data, seeds were targeted to each of the seven a priori selected networks in five cortical zones (rPFC, aINS, IPL, PCU, mPFC; see zones and labels in Fig. 1), (as shown in Fig. 2 and Supp. Figs. S3-S5). The one exception was default network A (DN-A) which does not include a region within the anterior insula, leaving four zones. In each participant (S1–S7, not including S2 who only provided a discovery dataset), seeds were defined in the discovery dataset, and used to calculate correlation values in the left-out, replication dataset. The larger matrix on the left shows the cross-correlation matrix averaged across all subjects, and the smaller matrices to the right show the correlations in each subject. In both the group-averaged and individual-subject matrices, clear separation of the PMN from other networks can be observed, with PMN-targeted seeds showing higher correlation with each other (diagonal; within-network correlations) than with other networks. Note that the PMN showed a weak negative correlation with DN-A and DN-B. The box plots show the group-averaged comparison between within-versus across-network correlations. For every network, within-network correlations were significantly higher than across-network correlations (t-tests, asterisks denote p< .05, Benjamini-Hochberg corrected for 6 pairs of network comparisons tested). This result was also significant when comparisons were performed within individuals (see Supp. Fig. S11) and in the left-out triplication data (Supp. Figs. S12-S13). This demonstrates that the PMN is separable from nearby networks to a similar degree as other networks are separated from each other, and that this dissociation is evident in multiple cortical zones. The top and bottom of each box represent the 75th and 25th percentiles, respectively, and the central mark within each box represents the median. Each dot represents a data point from each individual, the whiskers extend to the most extreme data points not considered outliers, and the outliers are plotted using dots beyond the whiskers. Default network-A (DN-A); default network-B (DN-B); frontoparietal control network A (FPN-A); frontoparietal control network B (FPN-B); salience network (SAL); cingulo-opercular network (CON).

### The PMN shows a repetition enhancement effect

A final set of analyses sought to confirm that the distributed PMN we identified using resting-state data showed the expected stimulus repetition enhancement effect reported in the literature (Gilmore et al., 2015, 2019; Nelson et al., 2013). We analyzed fMRI data from a continuous recognition task provided in the NSD during which the same participants viewed ∼10,000 images three times across sessions. First we plotted the task contrast maps comparing evoked responses for trials with repeated images (2^nd^ presentation, P2; 3^rd^ presentation, P3) to those with novel images (1^st^ presentation, P1). Fig. 8 demonstrates that the PMN, particularly in the posteromedial regions, exhibited a strong repetition enhancement effect in all individuals (Fig. 8). However, evidence for the effect was limited in other parts of the distributed PMN. We therefore conducted two targeted analyses to test whether the effect was observable in each cortical zone. First, we took the a priori defined seed vertices that were used to target networks in the resting-state data for statistical analysis (Supp. Figs. S3–S5), and calculated average trial-wise beta values for each session for each trial type (e.g., P1, P2, and P3). In all six individuals, the PMN exhibited a significant increase in activity (betas) as images were repeated (Supp. Fig. S16). This effect was observed in comparisons between P1 and P2 trial types and between P1 and P3 trial types (p < .001, Bonferroni corrected; Supp. Fig. S16, first row), but no significant differences were found between the P2 and P3 trials. Similar effects were observed in FPN-A, and DN-A showed the opposite effect, further supporting a functional separation between DN-A and PMN. Second, we used a regional approach and calculated the average betas across all vertices that were deemed to be part of the PMN by the clustering analysis within 5 broad cortical regions: the posterior midline (encompassing PCU, RMC and rPCC), anterior midline (encompassing mPFC), the posterior lateral cortex (encompassing IPL), anterior lateral cortex (encompassing rPFC), and the anterior insula. Because DN-A and DN-B showed evidence of negative correlation with PMN in the resting-state analysis (Fig. 7), we also calculated average regional beta values for these two additional networks. The matrices in Supp. Fig. S17 show that the PMN consistently showed a repetition enhancement effect in each of the 5 regions. In contrast, DN-A and DN-B tended to show a repetition suppression effect. Thus, the PMN showed a repetition enhancement effect in multiple cortical zones.

**Figure 8:**
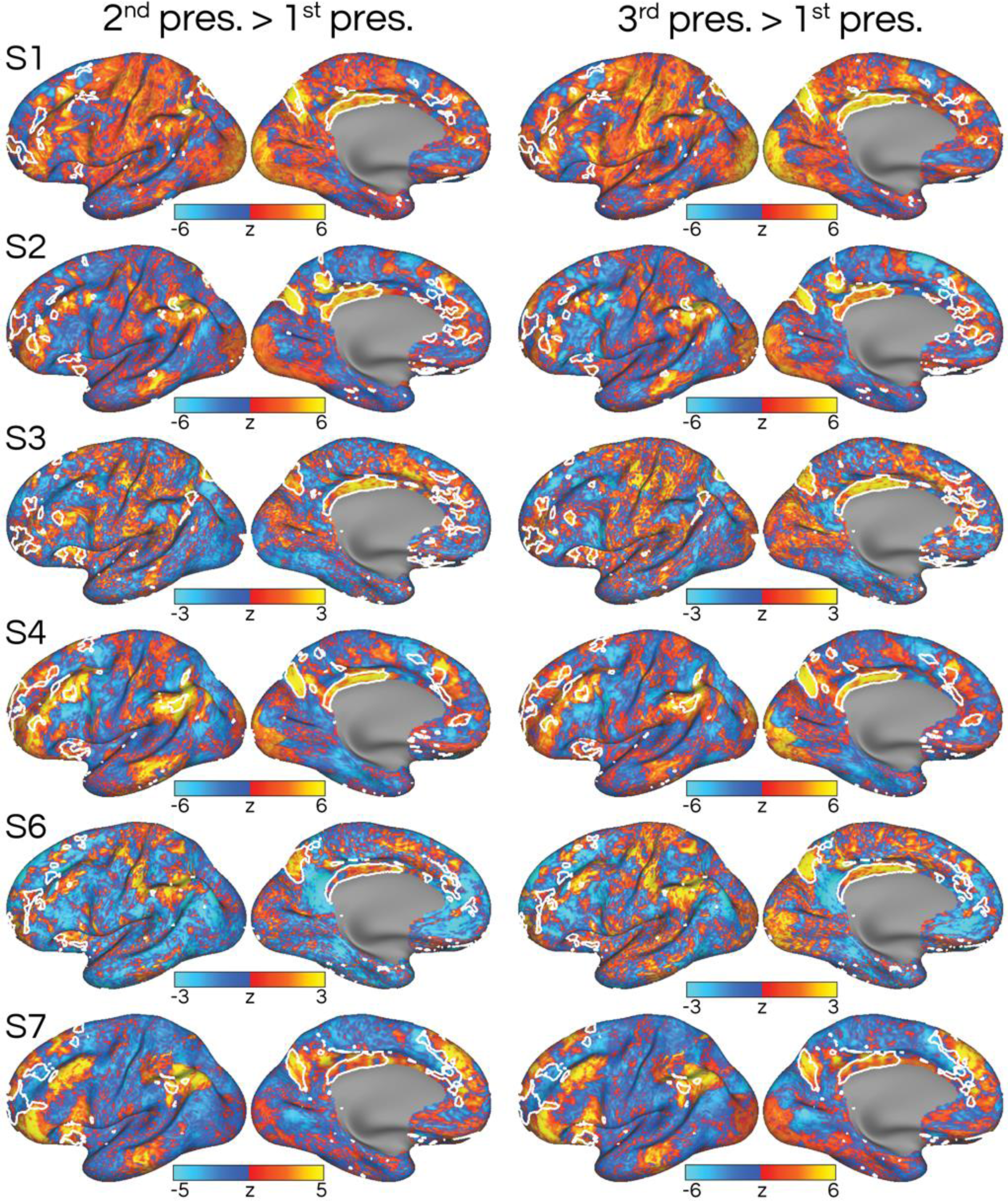
The estimated parietal memory network (PMN) shows a repetition enhancement effect. Estimates of trial-evoked activity were calculated for every trial of the Natural Scenes Dataset (NSD) experiment, in which subjects performed a continuous recognition task for naturalistic images. To determine if repeated presentations of the same image elicited higher activity (i.e., repetition enhancement), we contrasted beta estimates from trials in which images were shown a second or third time in each session of the task (i.e., “2^nd^ pres.” and “3^rd^ pres.”) against trials when the image was first presented (“1^st^ pres.”). Maps display z-scored *t* values representing the contrast between trial types, warmer colors indicate positive and cooler colors indicate negative values. The estimated boundaries of the PMN, derived from the resting-state clustering analysis (Fig. 1), are shown in white. The posteromedial regions of the PMN show the clearest repetition enhancement effect (warm colors), but evidence for the effect was also seen throughout the distributed extent of the network (Gilmore et al., 2015), and was confirmed by a region-of-interest analysis (Supp. Fig. S17).

## Discussion

We studied the detailed anatomy of the PMN using high-field and high-resolution 7T fMRI data. We found that, when defined within an individual, the PMN is a distributed network that contains regions in upwards of 9 cortical zones (Figs. 1 & 2), contrary to prior accounts that restricted the PMN to parietal and posteromedial cortices. We show that the PMN is closely interdigitated with, but clearly and statistically dissociable from, other nearby large-scale networks, including those within canonical default, frontoparietal control, and salience / cingulo-opercular network regions (Figs. 3–7 & Supp. Figs. S3–S13). The findings were confirmed in all 6 individuals analyzed (Fig. 1), were consistent across multiple analysis procedures (Figs. 1 & 2), were replicated and triplicated in the same individuals (Supp. Figs. S11–S14), were further replicated in an independent cohort of 8 individuals using a different scanner and sequence (Supp. Fig. S15), and were also confirmed in an analysis of data from 4,181 UK Biobank subjects (Miller et al., 2016; Supp. Fig. S19). These analyses all confirmed that the PMN is a distributed network with regions in multiple cortical zones outside of parietal and posteromedial cortex. The resulting organization suggests that the PMN presents as another instance of a parallel distributed network, an organization that is characteristic of association cortex (Goldman-Rakic, 1988) and likely emerges through the fractionation of a prototypical distributed network architecture during developmental processes of specialization (Buckner & Krienen, 2013; DiNicola & Buckner, 2021). Finally, we confirmed that the estimated PMN displays a repetition enhancement effect, even in these non-canonical regions such as the medial and lateral prefrontal cortex and anterior insula (Fig. 8 and Supp. Figs. S16 & S17).

### The PMN as a parallel distributed association network

In all individuals, the PMN was found to display a distributed organization, with regions in upwards of 9 cortical zones, including PCU, rPCC, RMC, mPFC, vmPFC, rPFC, dPFC, aINS, and IPL (see dashed boxes in Fig. 1). Sometimes these regions were small (e.g., see IPL and vmPFC in Fig. 1), and would have been overlooked if their clearer presence in other subjects were not suggestive. Fig. 1 shows the degree of overlap across individuals in the anatomical location of PMN regions. It is notable that the regions with the most overlap were also those that have most often been ascribed to the PMN: the PCU and rPCC (e.g., see Gilmore et al., 2015, 2021; Gordon et al., 2017). Other regions, such as the IPL and rPFC, showed more spatial variability across individuals, underscoring why they may have sometimes been missed. Similarly, although the rPFC and mPFC regions were relatively large, they were somewhat dispersed, leading to less overlap across individuals. One surprising finding was that the aINS showed high overlap across the 6 individuals tested in this high-resolution data, but is largely absent from prior estimates of the PMN (but see ‘Control C’ network in Kong et al., 2019). Because the insula is difficult to image due to the close positioning of opercular and insular cortices, this PMN region may have been missed due to blurring with nearby regions (see Supp. Fig. S10) in lower-resolution or group averaged approaches. The overlap map in Fig. 1 provides a compelling explanation for why different accounts of the PMN have been reported in group-averaged data.

An interesting observation was that half the individuals showed limited evidence of a further PMN region in the lateral temporal cortex. This would be unremarkable given small size of the region identified, the low correlation values, and its inconsistency across individuals. However, this putative region was located right next to a zone of signal dropout (Supp. Fig. S2), raising the possibility that a lateral temporal PMN region may also exist that has been missed. There are also reasons to think that this part of the brain should contain a PMN region. Our analyses suggest that the PMN is closely linked to FPN-A and FPN-B (Fig. 5 and Supp. Fig. S7), both of which contain prominently lateral temporal regions approximately where this putative PMN region might be (Fig. 3 and Supp. Fig. S15; and see Braga & Buckner, 2017; Doucet et al., 2011; Power et al., 2011; Yeo et al., 2011). Hence a lateral temporal PMN region might be predicted at this approximate location. Future work using techniques that further limit the signal dropout should verify the presence of this region. Notably, in our validation analysis of multi-echo fMRI data at 3T (which has larger voxels but theoretically reduces the effect of signal dropout) we observed a similar level (or lack) of evidence for this region.

### Relationship to other networks

Detailed analysis of the anatomy of PMN regions suggests that the PMN sits at the confluence of multiple networks, including DN-A, DN-B, FPN-A, FPN-B, CON, and SAL. At a fine scale, the PMN occupies regions that are often completely distinct from other nearby networks, despite the complex shape of regions involved (see detailed anatomy in Figs. 4–5 & Supp. Figs. S6–S9). Importantly, this organization was observed using both data-driven clustering and seed-based analyses of functional connectivity, the latter of which allows overlap as it does not enforce a winner-takes-all vertex assignment. Prior estimates have diverged in considering the PMN as a sub-system of the canonical default (e.g., Power et al., 2011) or frontoparietal control networks (e.g., Yeo et al., 2011). Here, in the posteromedial cortex (Fig. 4 and Supp. Figs. S6 & S10), three regions of the PMN at or near the PCU, rPCC, and RMC were found to encircle the regions of DN-A and DN-B. The prominence of the PCU and rPCC regions may have led to a stronger association in the literature between the PMN and the default network. However, the same rPCC region of the PMN is also juxtaposed with regions of FPN-A and FPN-B as one moves rostrally along the callosal sulcus (Fig. 3). Hence the PMN is closely juxtaposed next to frontoparietal control network regions in the posteromedial (Fig. 3), inferior parietal, and medial prefrontal cortices (Fig. 5). In the IPL, the PMN more often was juxtaposed with FPN-A and FPN-B, remarkably filling the small gap between the FPN-A and FPN-B in many individuals, and often not bordering DN-A or DN-B (Fig. 4). This variability, where the PMN borders DN-A and DN-B in some regions but not others, may be a result of higher variation in functional organization found in association cortex (Cui et al., 2020; Gao et al., 2014; Mueller et al., 2013; Qiu et al., 2022), or could be suggestive of further sub-structure within what we are defining as the PMN. Alternatively, these findings could suggest that the PMN is more closely linked to frontoparietal control network regions. Supporting this, the PMN was anti-correlated (i.e., showing negative correlations) with DN-A and DN-B (Fig. 7 & Supp. Fig. S12), but not FPN-A and FPN-B. These findings again support a separation of the functions of the PMN from those of DN-A and DN-B.

Particular consideration is given to the mPFC area, which here was defined as belonging to the PMN (based on connectivity to PCU and rPCC regions) but has in many accounts been ascribed to SAL (Gordon et al., 2017; Laumann et al., 2015; Schaefer et al., 2017; Zheng et al., 2022). In our analysis, we observed closely interdigitated mPFC regions of the PMN, SAL, and CON networks in two datasets (a total of 14 individuals) and confirmed the organization of the PMN in an analysis of UK Biobank data. The detailed anatomy shown in Supp. Figs. S8–S9 suggest that without high-resolution and individualized approaches these regions might easily be blurred together, which could lead to observations such as that SAL occupies the frontal components (aINS and mPFC) while the PMN occupies posterior components (PCU, rPCC and IPL) of this organization. Our analyses suggest that both networks include posterior and anterior regions that are closely interdigitated. To further confirm the distinction between the networks, we targeted seeds in an anterior-posterior progression along the mPFC to define PMN, SAL, CON, and a premotor network (PreM). This analysis was successful in 3 of the 6 NSD subjects in whom particularly good separation of networks was generally found. In each subject, the mPFC seeds defined distinct distributed networks (see white lines in Fig. 6). This confirmed the separation between PMN, SAL, and CON that we achieved in the main analyses, suggesting the three networks are distinct but very closely juxtaposed along the mPFC.

Overall, the PMN was found to be consistently positioned at the interface of approximately 5 large-scale networks (DN-A, DN-B, FPN-A, FPN-B, and SAL), with potentially a wider separation between PMN and CON. SAL regions were often positioned exactly between the two. The RMC regions of the PMN were closely juxtaposed with all 6 networks (Fig. 3 and Supp. Fig. S10). This positioning in between multiple networks was also evident in medial and lateral prefrontal cortex (Fig. 3), as well as the insula (Supp. Fig. S10). Despite the complex and detailed anatomy of juxtaposed regions, seeds targeted to each network in 5 cortical zones using the discovery dataset were statistically dissociated in independent data, both at the group (Fig. 7) and individual level (Supp. Fig. S11), and was replicated in the triplication dataset (Supp. Figs. S12 & S13). These results indicate that the PMN is as distinct from other large-scale networks as they are distinct from each other, with higher in-network (r ∼ 0.5) than across-network (r < 0.25) correlations (Fig. 7).

### Functional dissociation from canonical default regions

In line with previous studies, analysis of a continuous recognition task provided further evidence for the separation between PMN and canonical default network regions. We observed the repetition enhancement effect robustly within the PMN (Fig. 8), but observed the opposite effect in DN-A (Fox et al., 2005). Notably, DN-A and DN-B showed evidence of being anticorrelated with PMN in Fig. 7 and Supp. Fig. S12. In addition, we also observed a repetition enhancement effect in FPN-A and FPN-B (Supp. Fig. S16). These results, along with the close juxtaposition between PMN and frontoparietal control networks in regions such as the IPL, mPFC and aINS, suggest that the PMN may be more closely aligned functionally to the frontoparietal control than default network systems.

### Limitations and technical considerations

Although care was taken to ensure that the networks were accurately identified and were consistent across individuals and estimation methods, it is possible that in some cases our clustering analyses over-split certain networks. For instance, in the case of three individuals (S1, S6 & S7), the clustering solution (i.e., value of *k*) used led to a division of the canonical default network into three networks, rather than two as per our previous estimates (Braga & Buckner, 2017). In these individuals, we carefully compared the clustering-based networks to our seed-based estimates and to maps from task-based analyses targeting DN-B and DN-A in the same individuals (data not shown) and ascertained that the three-network clustering solution was likely due to over-splitting. In these cases, we took the two networks that were closest to the PMN and labelled them “DN-A” and “DN-B”, however the results should be interpreted accordingly: these subjects were missing some components of DN-A (see descriptions in Braga et al., 2019; Braga & Buckner, 2017) such as the key region that extends into ventral posterior cingulate and retrosplenial cortex (see Fig. 3). Hence in these participants “DN-A” should be considered with this caveat. Note this does not affect the claims about the PMN being distinct and distributed. This over-splitting was not present in the three subjects that we focus on more extensively (S2, S3 & S4; Figures 4 & 5 and Supp. Fig. S8). The analyses here also focus heavily on resting-state functional connectivity, and need to be supported by task-based analysis within extensively sampled individuals, at high-resolution, to further confirm the distinctions we report, beyond the task reported here. Finally, here we focused on the cerebral cortex, but it is likely that the PMN includes additional regions, such as those reported in the posterior medial temporal lobe (Zheng et al., 2021), which may play a crucial role in understanding the function of the PMN and how it relates to other networks.

## Conclusion

In summary, here we show that the PMN is a distinct, distributed network with regions in upwards of 9 cortical zones. We show that the PMN is closely juxtaposed with approximately 6 large-scale networks (DN-A, DN-B, FPN-A, FPN-B, SAL, CON), with some evidence for a closer link between the PMN and frontoparietal control regions based on spatial proximity and similarity of task-evoked responses. The results underscore the need for individualized, high-resolution, and high-field fMRI studies that provide greater separation between the small and tightly interwoven regions that populate the cortical mantle.

## Acknowledgements

This research was supported in part through the computational resources and staff contributions provided for the Quest high performance computing facility at Northwestern University which is jointly supported by the Office of the Provost, the Office for Research, and Northwestern University Information Technology. This work was supported by the Center for Translational Imaging at Northwestern University.

---

**Supplementary Figure S1:**
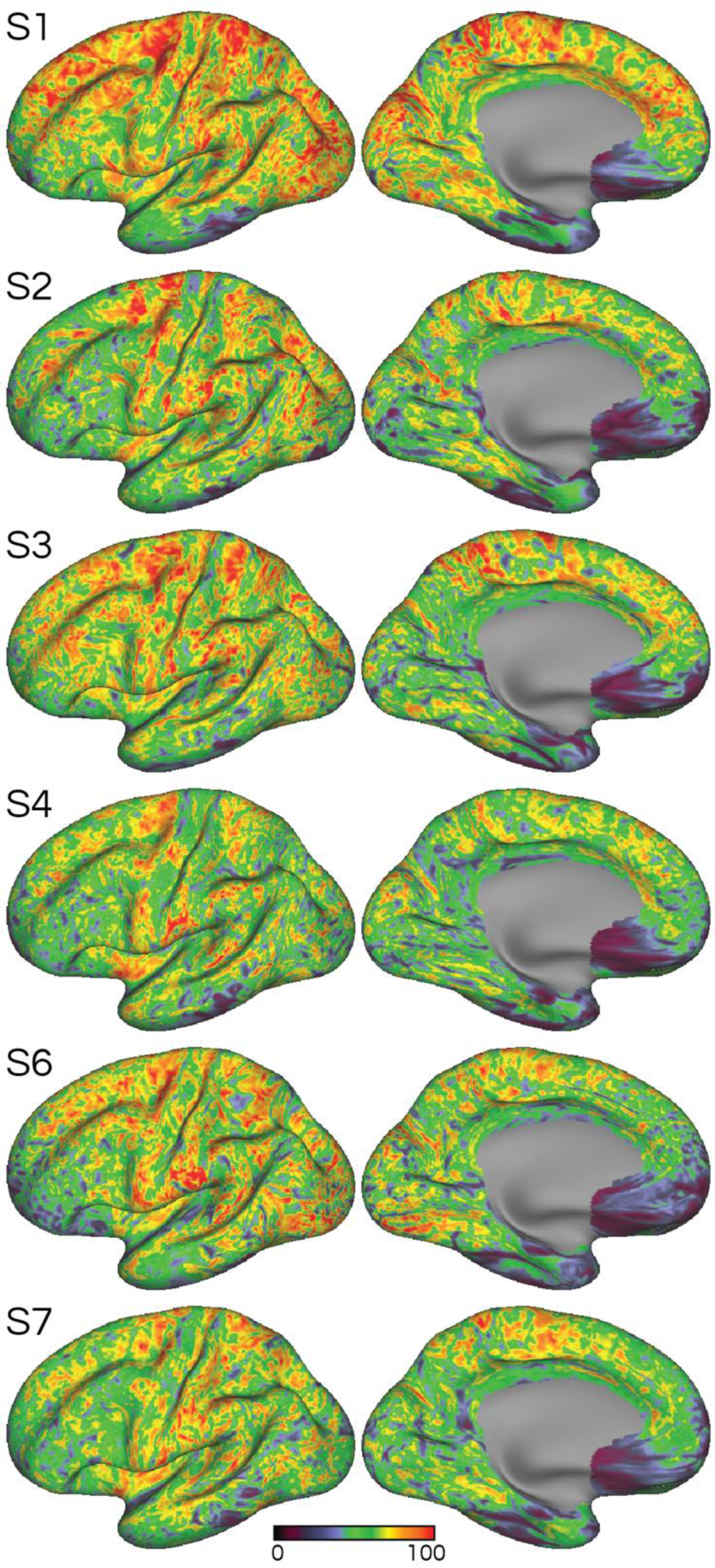
Good coverage and high signal-to-noise ratio was achieved in the high-resolution 7T data. Temporal signal to noise ratio maps for all six individuals (S1-S7; rows). The maps show that good data quality and coverage was achieved by the high-field 7T protocol, despite the small voxel size (1.8 mm isotropic). Signal dropout regions (cooler colors) can be seen in the temporal pole, lateral temporal cortex, and ventromedial prefrontal cortex.

**Supplementary Figure S2:**
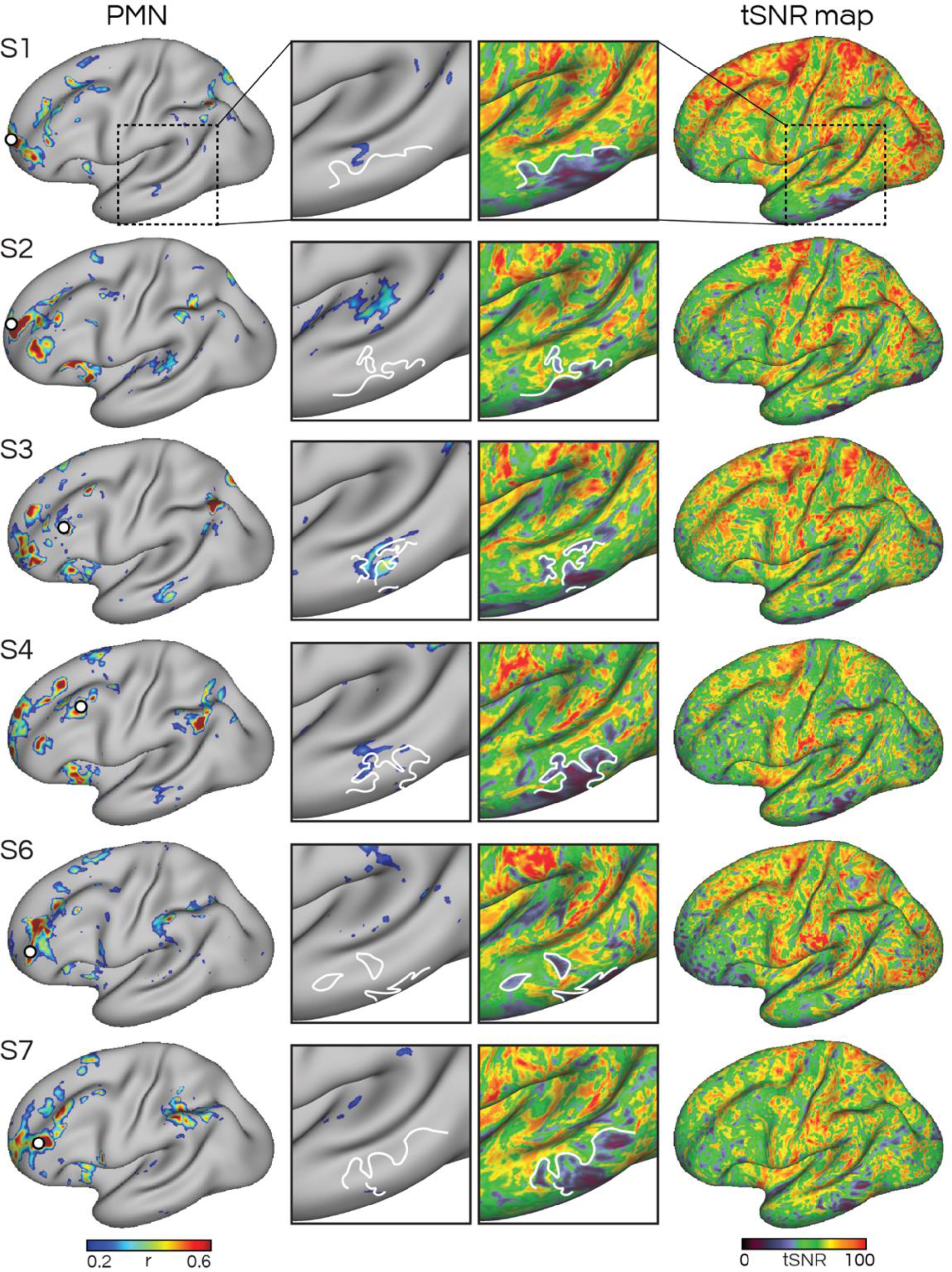
Signal dropout may have affected detection of a PMN region in the lateral temporal cortex. The left column shows the functional connectivity map of the PMN defined using a seed-based approach (seeds shown as white circles), and the right column shows the temporal signal-to-noise ratio (tSNR) maps. Insets show a zoom-in of the lateral temporal cortex, focused on signal dropout regions in each individual. The white lines trace the idiosyncratic shape of regions affected by signal dropout in each individual. In 3 subjects (S1, S3, S4), evidence for a region of the PMN was detected in close proximity to signal dropout regions, raising the prospect that a lateral temporal PMN region may have been missed in other participants. Analysis of UK Biobank data containing 4,181 participants also suggested the presence of a PMN region here (see Supp. Fig. S19).

**Supplementary Figure S3:**
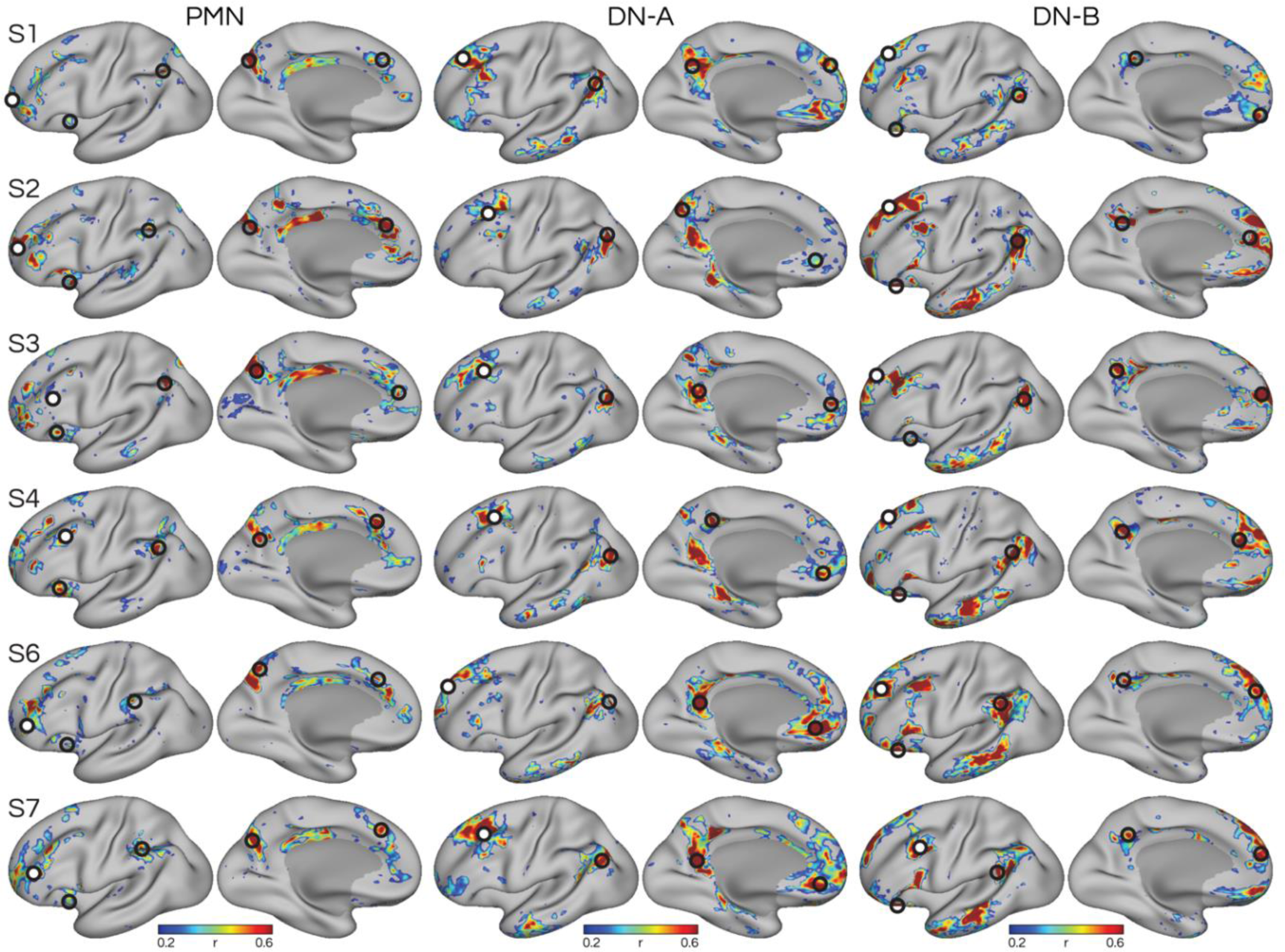
Seed-based network estimation confirms the parietal memory network (PMN) is distinct from nearby distributed networks within the canonical default network. Seeds were selected from the rostral prefrontal cortex (white circles) in each individual (S1–S7; rows) targeting three distributed networks: the PMN, default network A (DN-A), and default network B (DN-B; Braga and Buckner 2017). Black hollow circles represent seeds targeting regions of each network in four other cortical zones (anterior insula, inferior parietal lobule, precuneus, medial prefrontal cortex), with the exception of DN-A which did not display an anterior insula region. These seeds were used to provide converging estimates of the networks, and for statistical analyses in left-out data. The three networks contained regions that were closely juxtaposed in multiple cortical zones but occupying distinct parts of the cortex (see detailed anatomy in Fig. 4 and Supp. Fig. S6). Thus, despite all three networks containing prominent posteromedial regions, each could be clearly separated through within-individual resting-state functional connectivity at high resolution.

**Supplementary Figure S4:**
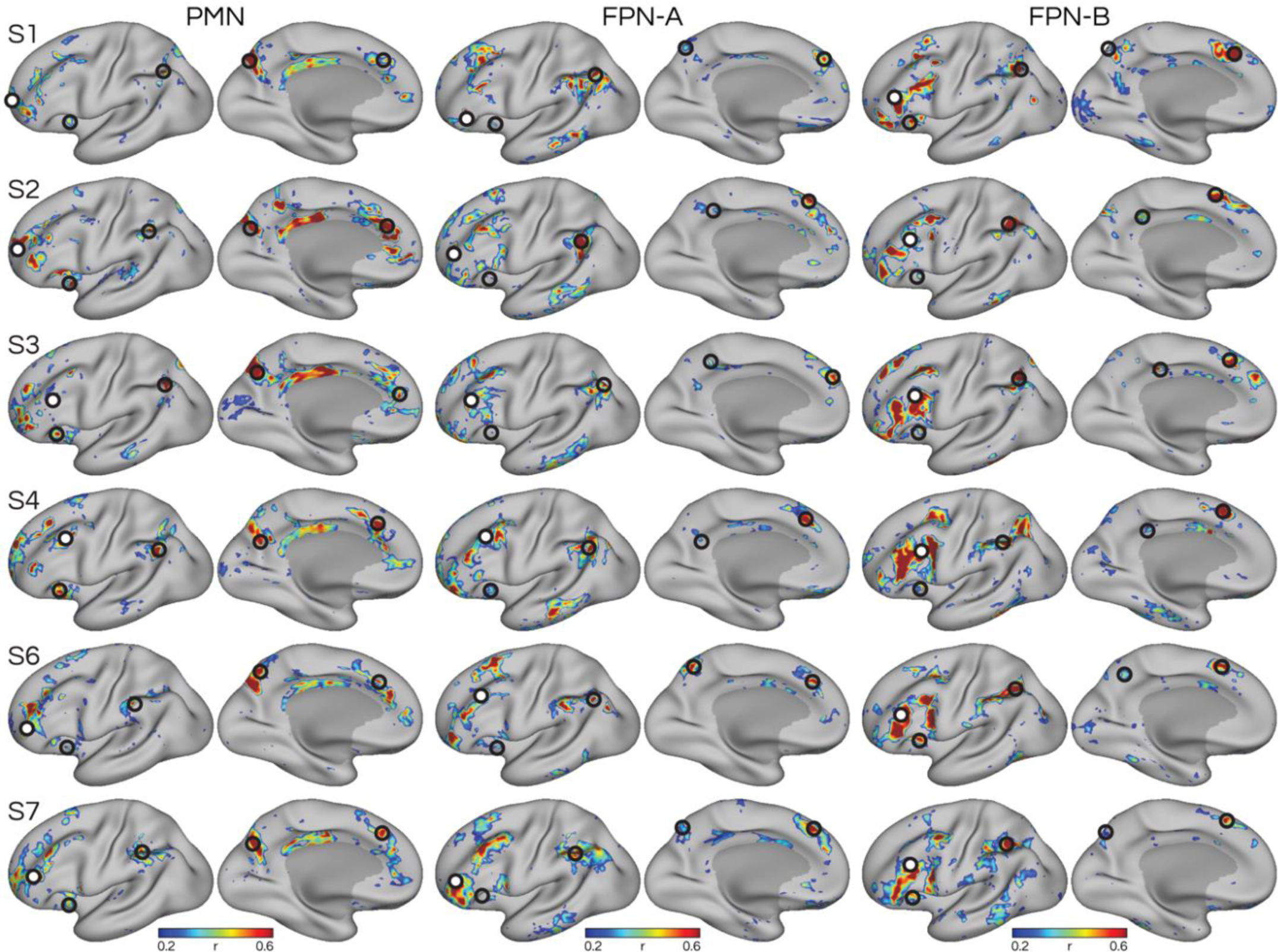
Seed-based network estimation confirms the parietal memory network (PMN) is distinct from nearby distributed networks within the canonical frontoparietal control network. Figure is formatted according to Supp. Fig. S3. Seeds were selected from the prefrontal cortex (white circles) in each individual (S1–S7; rows) targeting three distributed networks: the PMN, frontoparietal network A (FPN-A), and frontoparietal network B (FPN-B; Braga and Buckner 2017). The three networks contained regions that were closely juxtaposed in multiple cortical zones but with each network occupying distinct parts of the cortex (see detailed anatomy in Fig. 5 and Supp. Fig. S7).

**Supplementary Figure S5:**
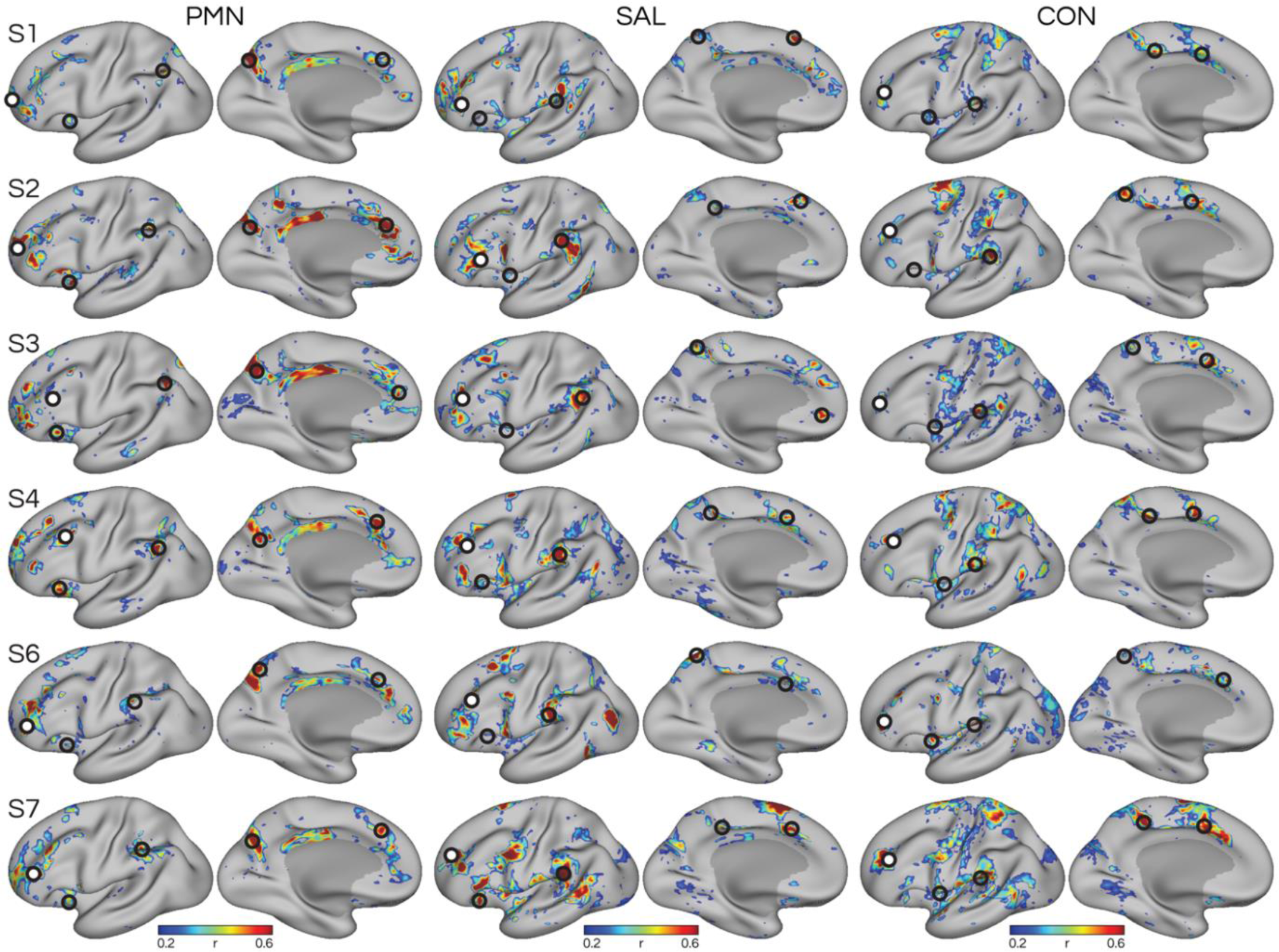
Seed-based network estimation confirms the parietal memory network (PMN) is distinct from nearby distributed networks within the canonical salience/cingulo-opercular network. Figure is formatted according to Supp. Fig. S3. Seeds were selected from the prefrontal cortex (white circles) in each individual (S1–S7; rows) targeting three distributed networks: the PMN, salience network (SAL), and cingulo-opercular network (CON). The three networks contained regions that were closely juxtaposed in multiple cortical zones but with each network occupying distinct parts of the cortex (see detailed anatomical distinctions in Supp. Figs. S8–S9).

**Supplementary Figure S6:**
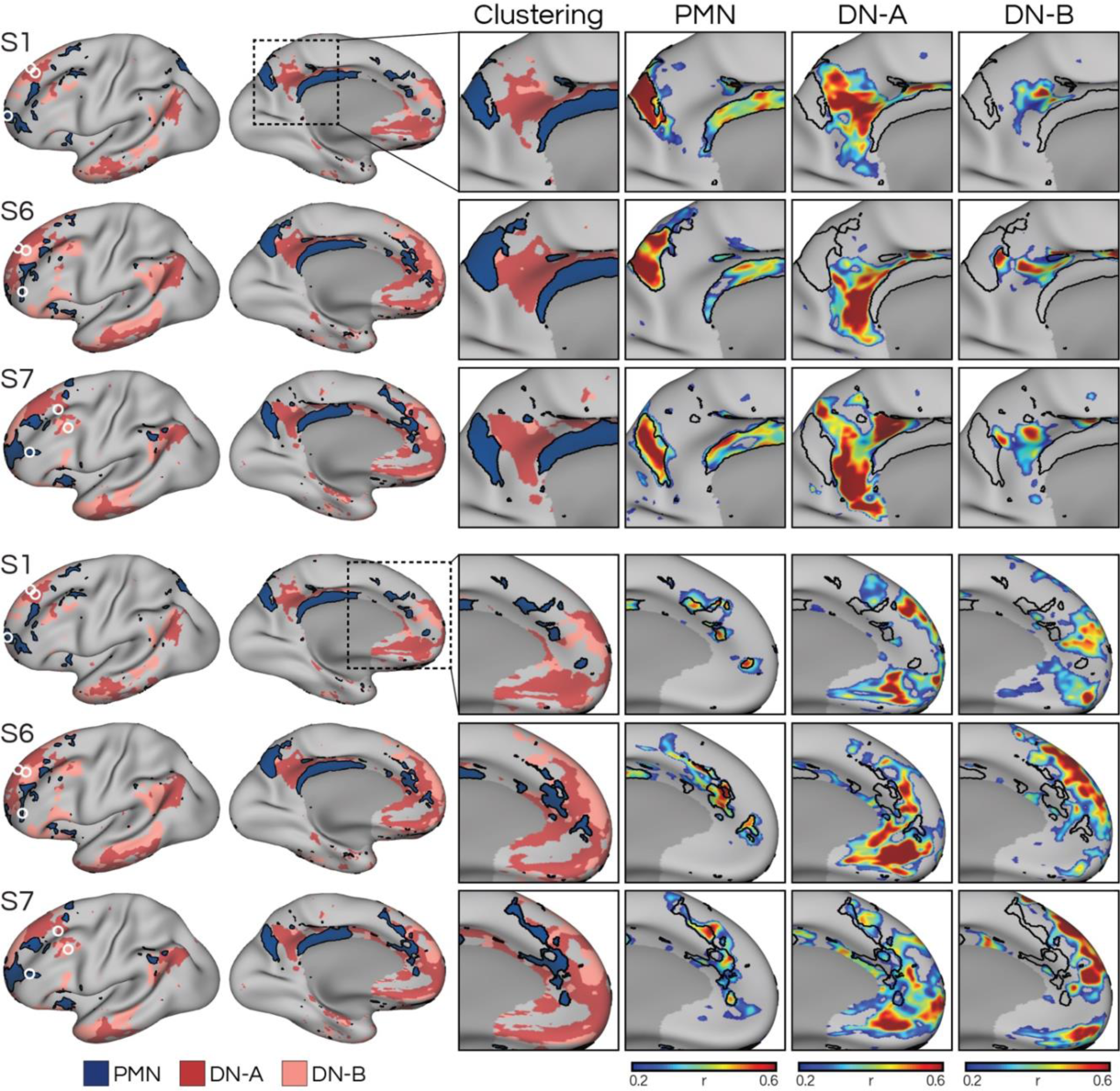
Detailed anatomy of the parietal memory network (PMN) reveals regions that are closely-knit with but distinct from networks within the canonical default network (DN) in additional individuals. Figure formatted according to Fig. 4. Insets show a zoom-in of the posteromedial (top three rows) and medial prefrontal (bottom rows) cortex, showcasing detailed anatomical distinctions between network regions. The black lines represent the boundaries of the PMN calculated from the parcellation approach, to serve as landmarks. Three other individuals (S2, S3, and S4) are shown in Fig. 4.

**Supplementary Figure S7:**
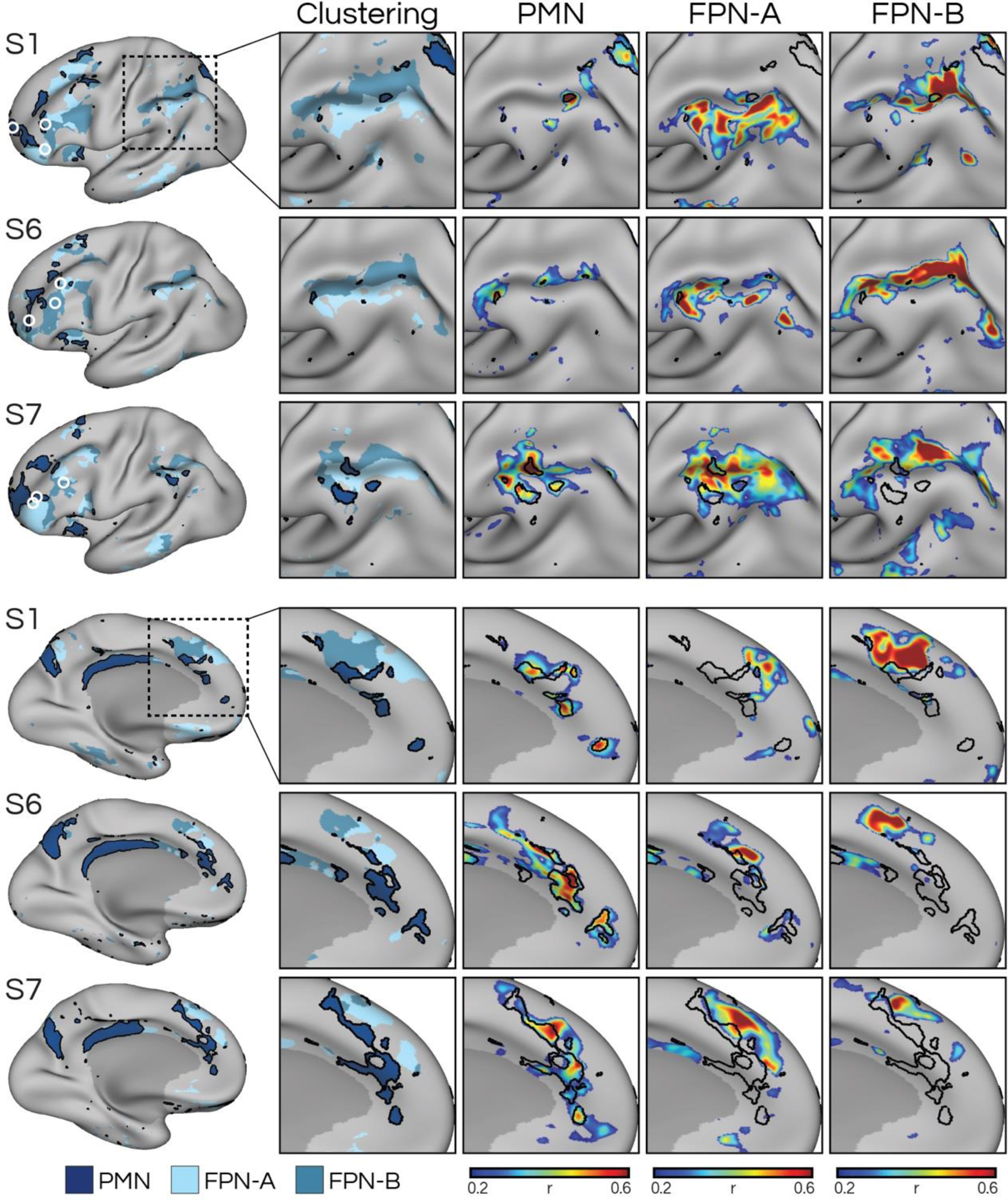
Detailed anatomy of the parietal memory network (PMN) reveals regions that are closely-knit with but distinct from networks within the canonical frontoparietal control network (FPN) in additional individuals. Figure formatted according to Fig. 5. Insets show a zoom-in of the intraparietal sulcus (rotated; top three rows) and anterior midline (bottom rows), showcasing the detailed anatomical distinctions between regions. In each cortical zone, the PMN occupied distinct regions that were closely juxtaposed with regions of frontoparietal control network A (FPN-A) and B (FPN-B). Three other individuals (S2, S3, and S4) are shown in Fig. 5.

**Supplementary Figure S8:**
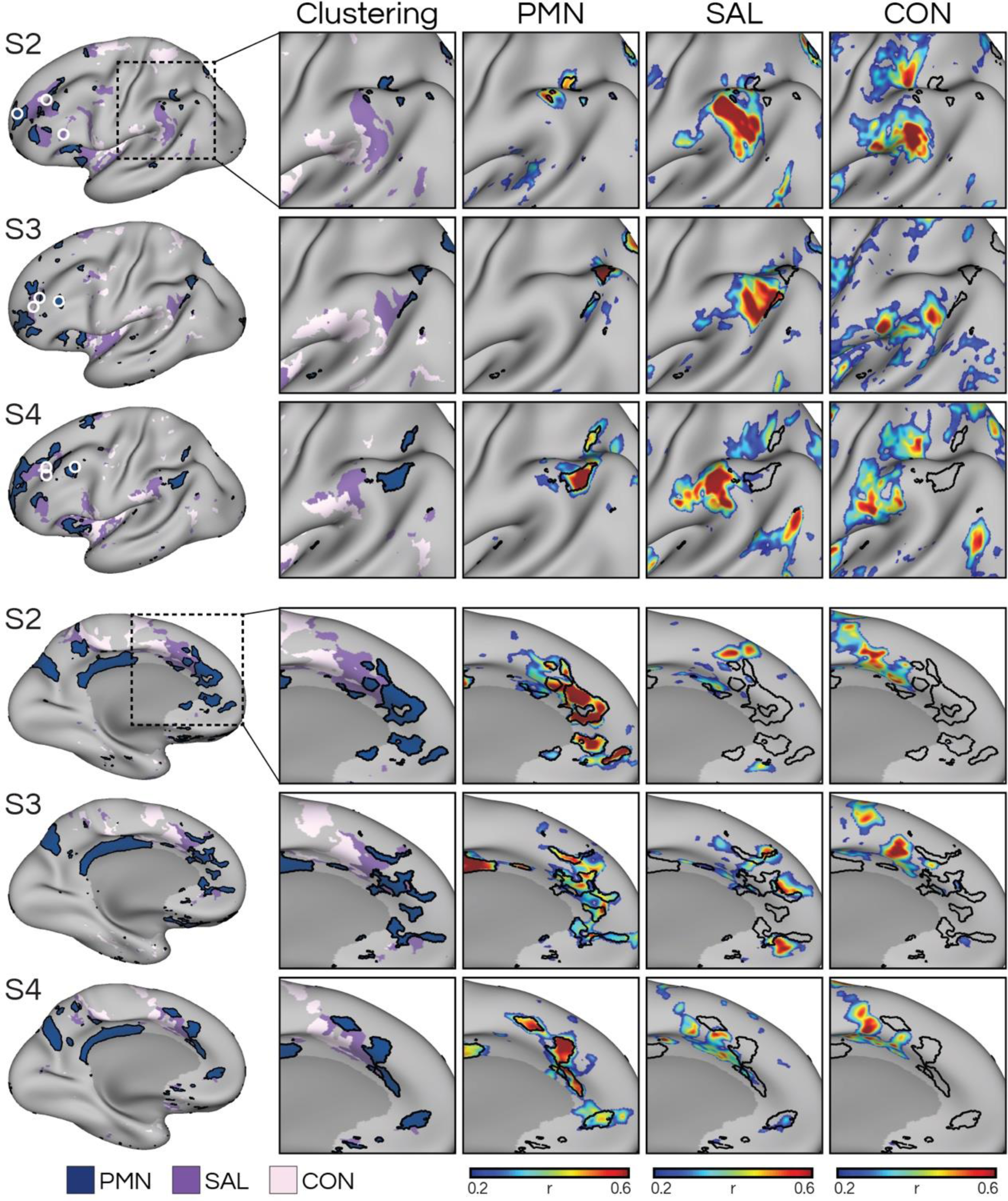
Detailed anatomy of the parietal memory network (PMN) reveals regions that are closely-knit but distinct from the salience (SAL) and cingulo-opercular (CON) networks. Left column shows the clustering-defined networks from Fig. 3 and the location of manually selected seeds (white circles) initially used to define the networks (see Supp. Fig. S5). Insets show a zoom-in of the intraparietal sulcus (top three rows) and medial prefrontal cortex (bottom rows), showcasing the interdigitation of network regions. The PMN occupied distinct regions that were closely juxtaposed with regions of the salience and cingulo-opercular networks. We also performed a targeted analysis of the distinction between the three networks in Fig. 6. Three representative participants (S2, S3, and S4) are shown, with the remaining 3 shown in Supplementary Figure S9.

**Supplementary Figure S9:**
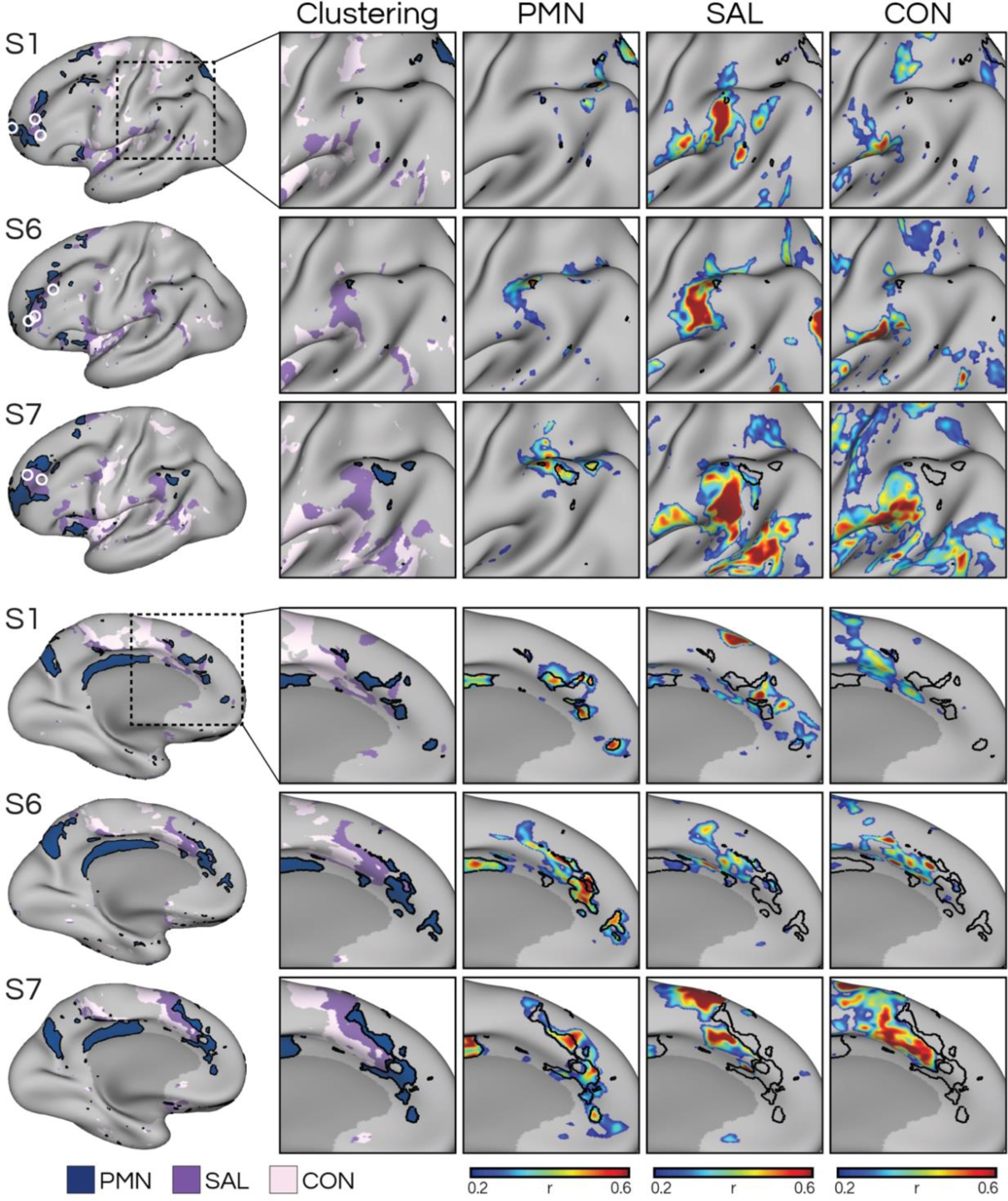
Detailed anatomy of the parietal memory network (PMN) reveals regions that are closely-knit with but distinct from the salience (SAL) and cingulo-opercular (CON) networks in additional individuals. Figure formatted according to Supp. Fig. S8. Insets show a zoom-in of the intraparietal sulcus (rotated; top three rows) and anterior midline (bottom rows), showcasing the detailed anatomical distinctions between regions. In each cortical zone, the PMN occupied distinct regions that were closely juxtaposed with regions of the salience and cingulo-opercular networks. Three other individuals (S2, S3, and S4) are shown in Supp. Fig. S8. See also targeted analysis of these networks in Fig. 6.

**Supplementary Figure S10:**
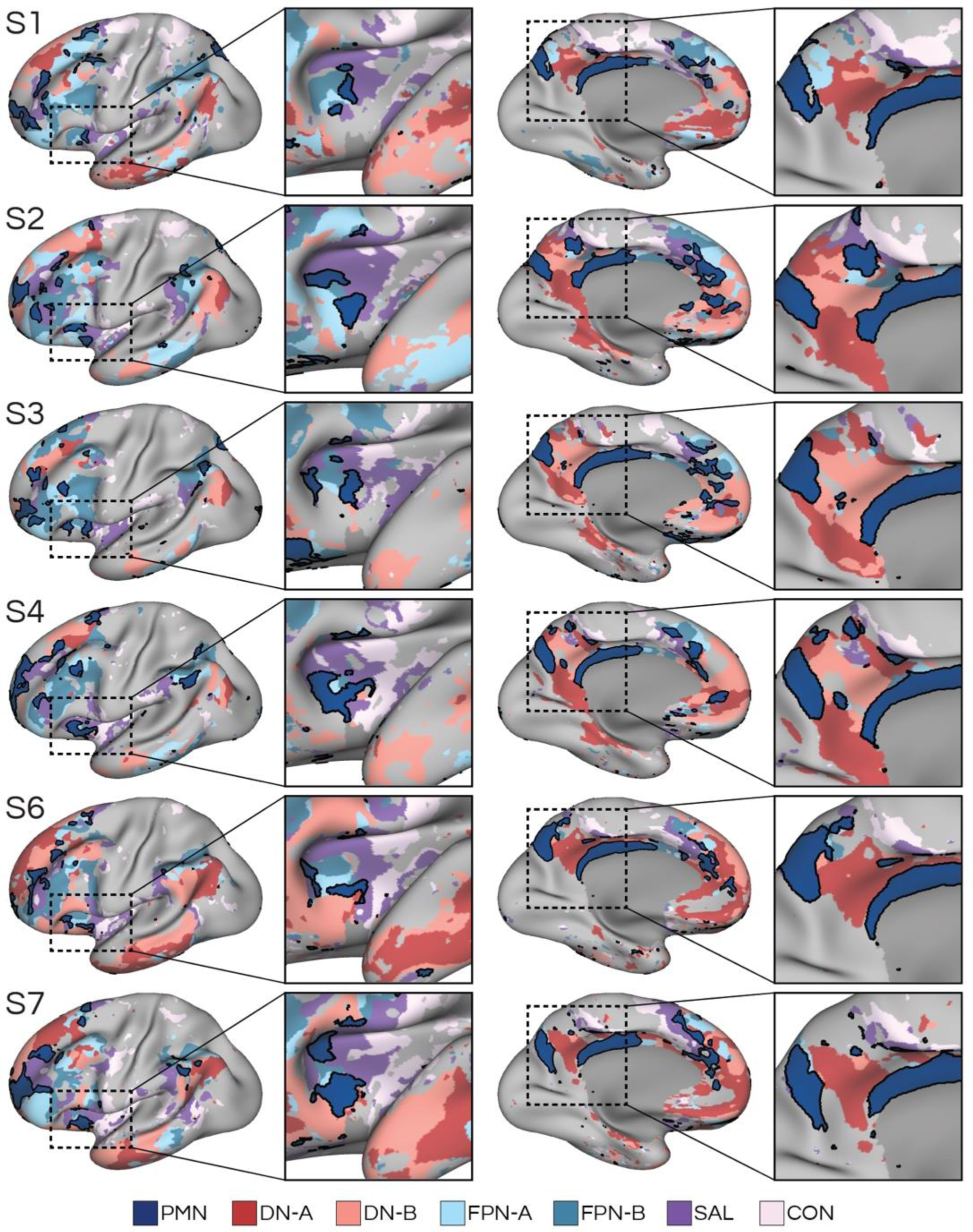
Detailed anatomy of networks within the posteromedial cortex and anterior insula reveals closely-knit organization of networks. The PMN was consistently positioned at the interface of six distributed networks. In the posteromedial cortex, three PMN regions, at or near the precuneus, rostral posterior cingulate, and ramus marginalis, were found to encircle the regions of DN-A and DN-B. In the anterior insula, the PMN was situated between FPN-A, FPN-B, SAL, and CON.

**Supplementary Figure S11:**
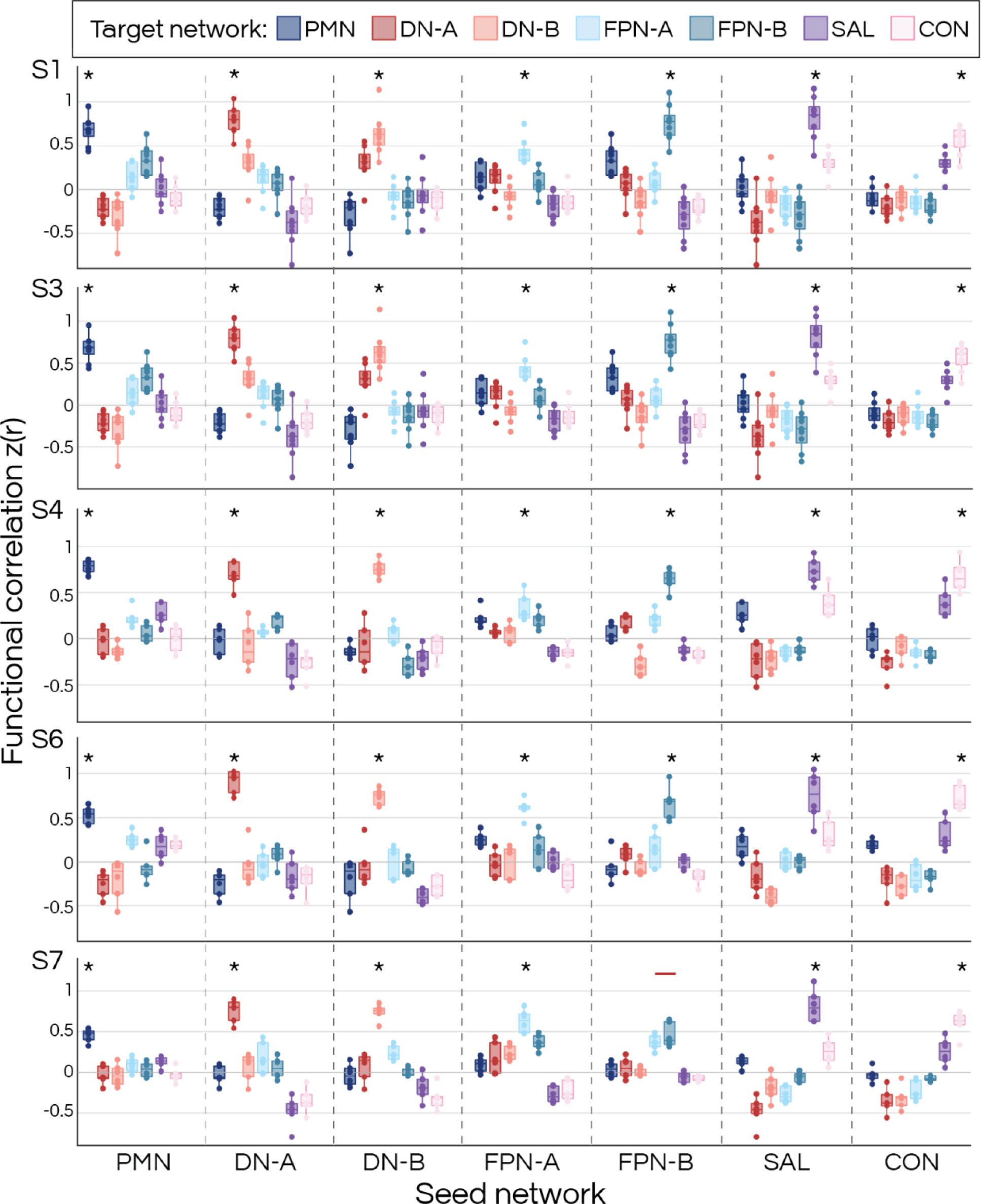
Within-individual statistical dissociation of the parietal memory network (PMN) from nearby networks in left-out replication data. The box plots show the comparison between within-versus across-network functional connectivity (r) for each individual (rows) for each network (columns divided by dashed lines). Asterisks indicate that the network’s within-network correlations are statistically higher than across-network correlations (* p<.05, Benjamini-Hochberg corrected for six pairs of network comparisons tested). Red lines indicate that the difference between the pairs are not significant. This demonstrates that the PMN is separable from nearby networks to a similar degree as other networks are separated from each other, and that this dissociation is evident in multiple cortical zones. Box plots formatted according to Fig. 7.

**Supplementary Figure S12:**
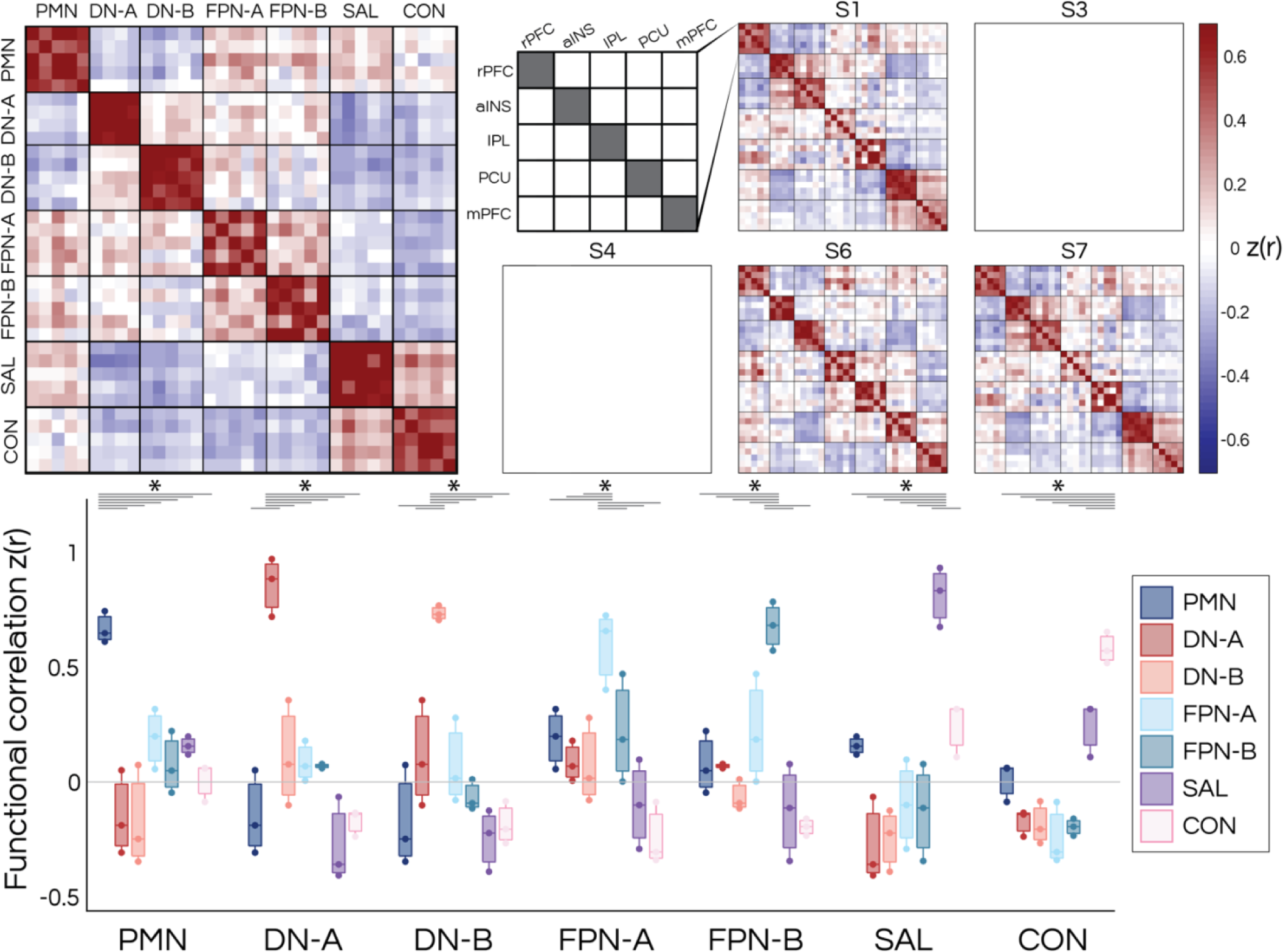
Statistical dissociation of the parietal memory network (PMN) from nearby networks in left-out triplication data. The dissociation of the PMN from nearby networks in left-out data was replicated in individuals who provided sufficient data for a triplication dataset (S1, S6, S7). Figure formatted according to Fig. 7. This demonstrates that the PMN is separable from nearby networks to a similar degree as other networks are separated from each other, and that this dissociation is evident in multiple cortical zones.

**Supplementary Figure S13:**
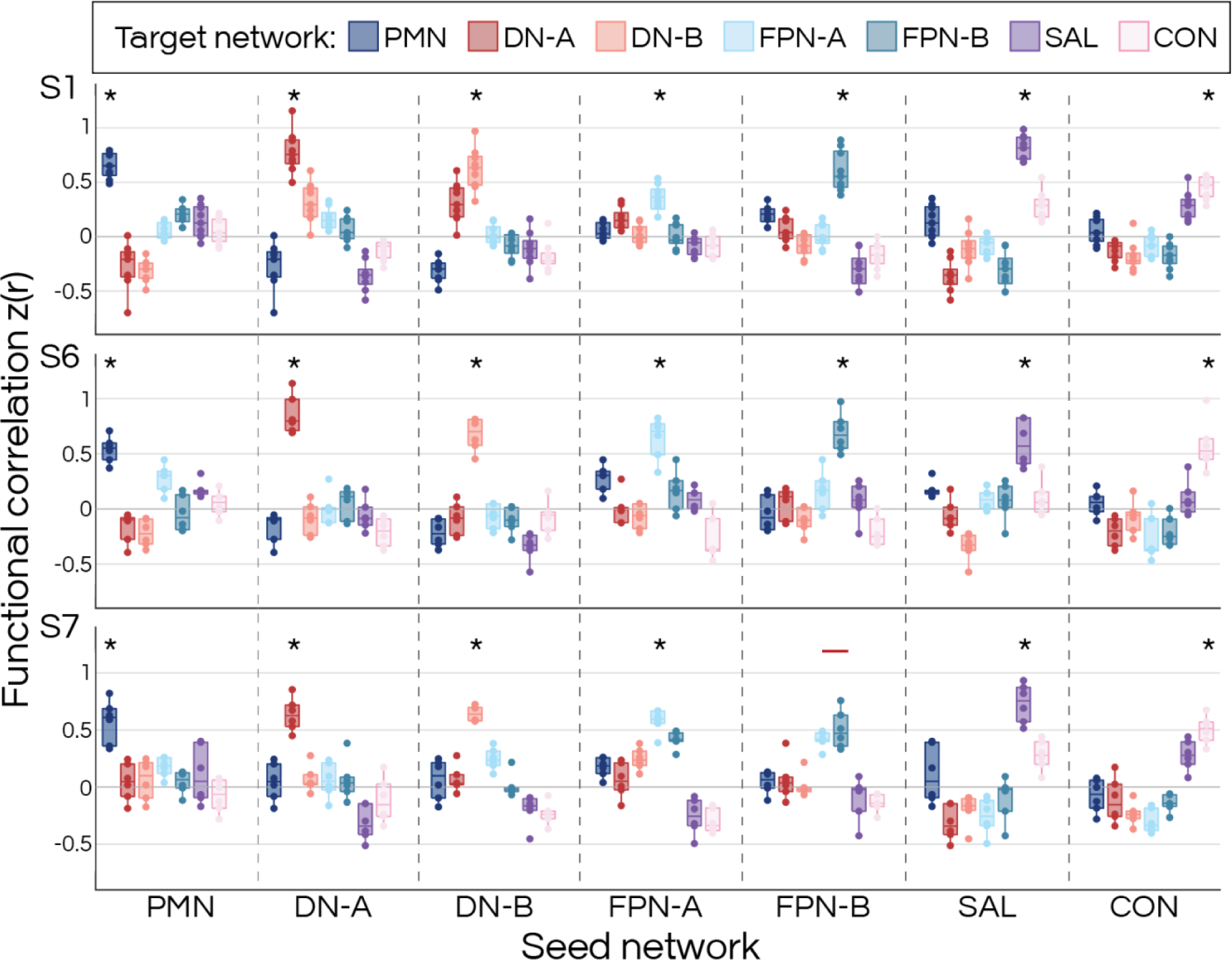
Within-individual statistical dissociation of the parietal memory network (PMN) from nearby networks in left-out triplication data. Figure formatted according to Supp. Fig. S11. The box plots show the comparison between within-versus across-network functional connectivity (r) for each individual who provided sufficient data for a triplication dataset (S1, S6, S7; rows) for each network (columns divided by dashed lines). This confirms that the PMN is separable from nearby networks to a similar degree as other networks are separated from each other, and that this dissociation is evident in multiple cortical zones.

**Supplementary Figure S14:**
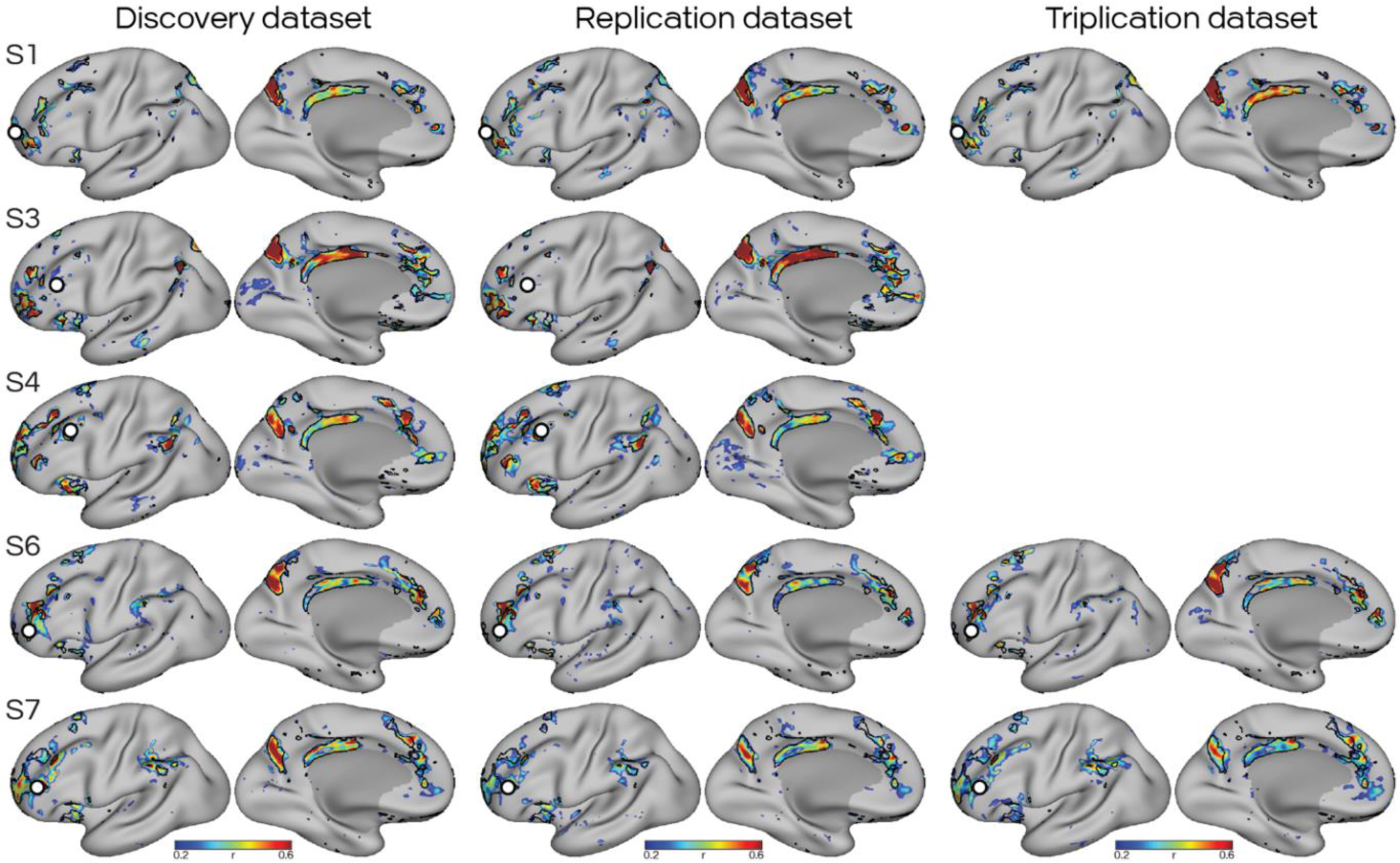
The distributed organization of the parietal memory network (PMN) is replicated and triplicated in the left-out data. High-quality resting-state runs from each participant were divided into a discovery dataset and 1 or 2 validation (i.e., replication and triplication) datasets, depending on data availability. Once statistical analysis had been run in the left-out validation data (Fig. 7 and Supp. Figs. S11-S13), the seeds selected from the discovery dataset (white circles) were applied to the two validation datasets. The black lines represent the boundaries of the PMN calculated from the clustering approach using the discovery dataset. The converging estimates demonstrate that PMN reliably includes regions distributed throughout multiple cortical zones.

**Supplementary Figure S15:**
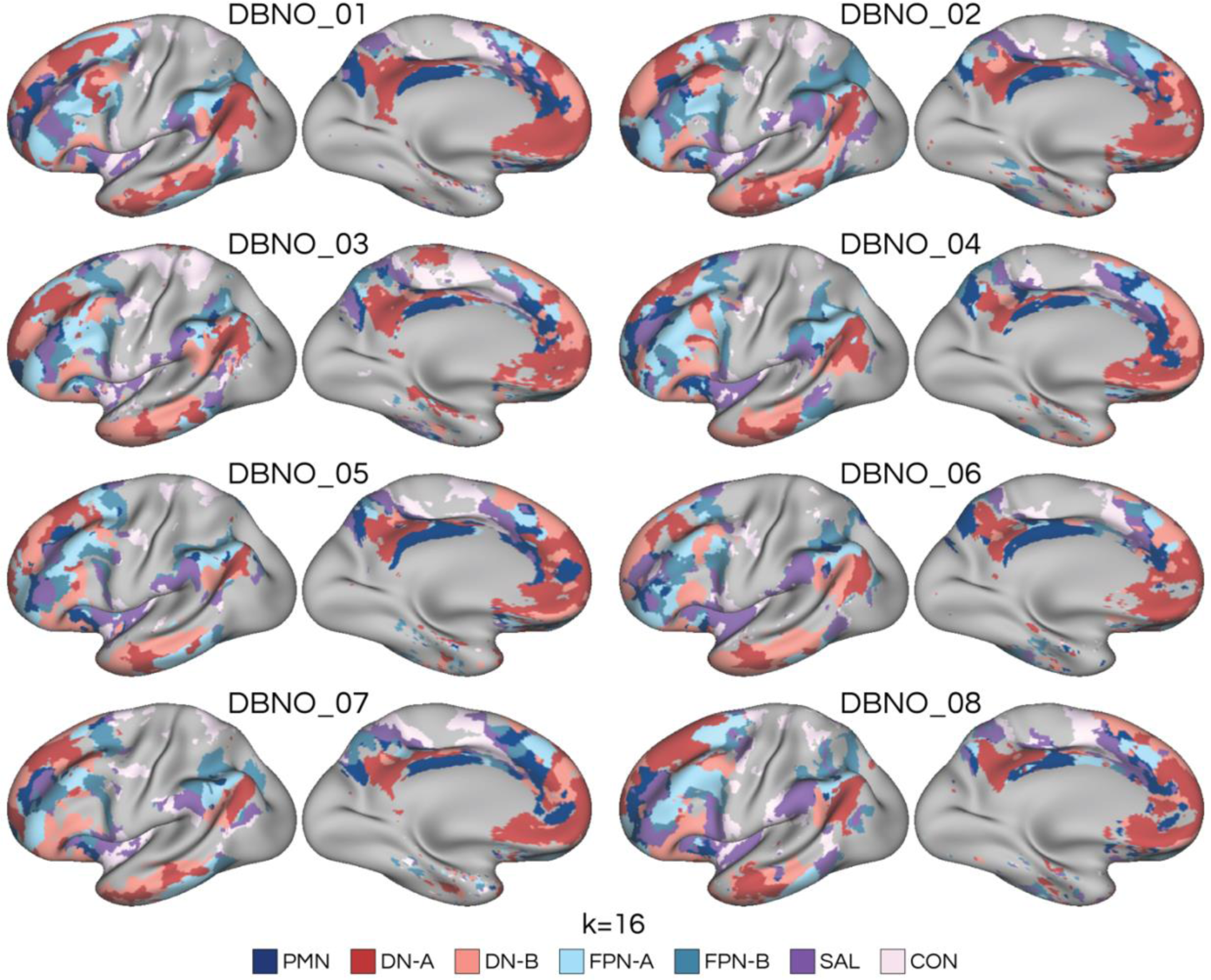
Replication of the parietal memory network (PMN) as a distributed network in an independent dataset of 8 additional individuals. Functional connectivity of 7 a priori networks were estimated using a clustering approach. Networks were selected from the clustering solution based on anatomical distinctions reported in the literature (Braga and Buckner 2017; Braga et al. 2019; 2020; Kong et al., 2019; Yeo et al., 2011). The independent ‘detailed brain network organization’ (DBNO) dataset replicated, within additional individuals at 3T, and using a different scanner and protocol, that the PMN is a distributed network. The clustering solution using k=16 from each individual is shown, representing the lowest value of k that separated the full set of a priori selected networks in each subject.

**Supplementary Figure S16:**
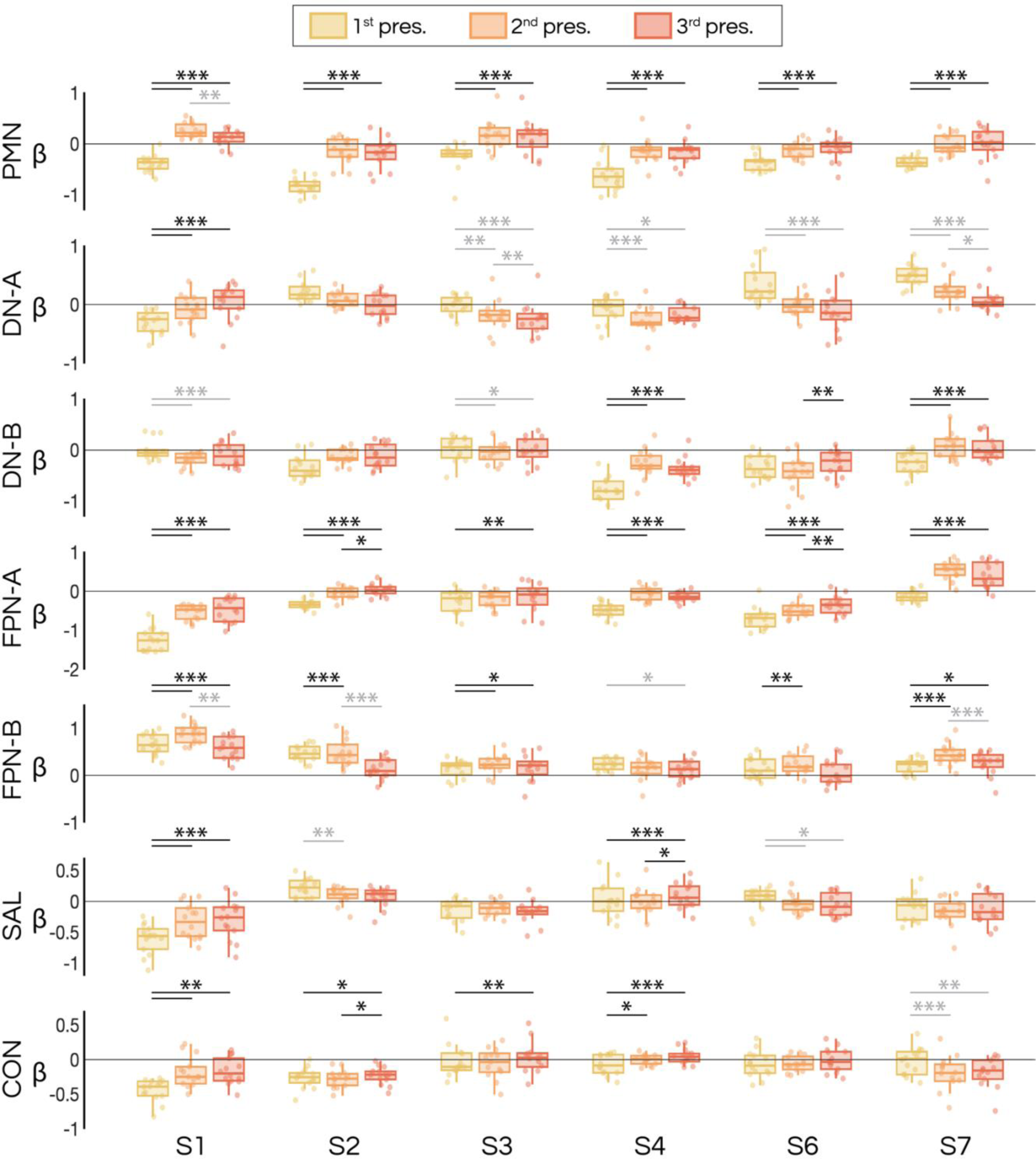
The parietal memory network (PMN) shows a repetition enhancement effect in all individuals. Box plots show the averaged estimates of the trial-evoked activity for 1^st^, 2^nd^, and 3^rd^ presentations of images in the continuous recognition task. The mean betas were calculated for each network by averaging betas from the seeds targeted to each network within five cortical zones, as used in the resting-state functional analysis (Fig. 7). The PMN showed a consistent increase (i.e., the repetition enhancement effect) in all individuals (black bars comparing 2^nd^ and 3^rd^ presentations to 1^st^). Other networks, particularly FPN-A, also showed evidence of statistically significant increases (black bars) and decreases (gray bars) in some individuals. A t-test was used to compare trial types (*p < .05; **p < .01; ***p < .001, Bonferroni corrected for the 3 pairs of trial types). Each data point represents the averaged beta from each session the whiskers extend to the most extreme data points not considered outliers, and the outliers are plotted using dots beyond the whiskers.

**Supplementary Figure S17:**
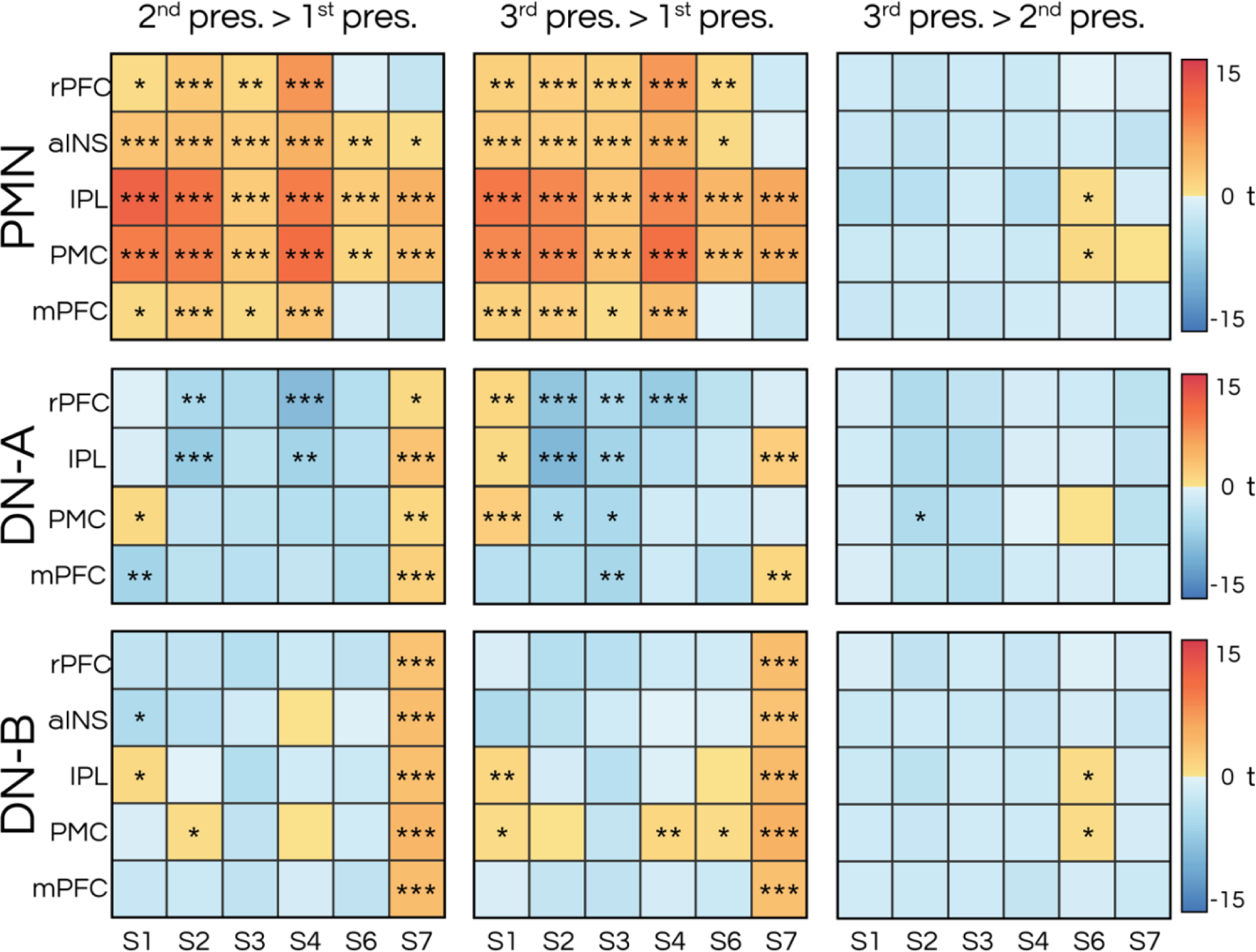
The parietal memory network (PMN) shows a repetition enhancement effect across multiple network regions, which was not observed in default network A (DN-A) and B (DN-B). The PMN shows increased activation when images are presented for the 2^nd^ and 3^rd^ time. In contrast, DN-A and DN-B show a tendency to deactivate when images are repeated. Boxes show the difference in trial-evoked activity (betas) between 1^st^, 2^nd^, and 3^rd^ appearance of images in the continuous recognition task. Each row corresponds to a different network (PMN, DN-A, and DN-B), and each column represents the comparisons between the trial types. Within each matrix, the rows represent five regions, including the anterior lateral cortex (encompassing rPFC), the anterior insula, the posterior lateral cortex (encompassing IPL), posteromedial cortex (PMC; encompassing PCU, RMC and rPCC), and anterior midline (encompassing mPFC), and each column denotes a different individual (S1-S7). The mean betas for each region were calculated by averaging all vertices within network regions defined through the clustering approach using resting-state data (shown in Fig. 3). A t-test was performed to compare trial types (*p < .05; **p < .01; ***p < .001, Bonferroni corrected for the 3 pairs of trial types). In all individuals, the PMN shows an increase in activation (i.e., the repetition enhancement effect, warmer color) in all or a majority of regions, when comparing the 2^nd^ and 3^rd^ presentation to 1^st^. This was not consistently observed for the DN-A and DN-B.

**Supplementary Figure S18:**
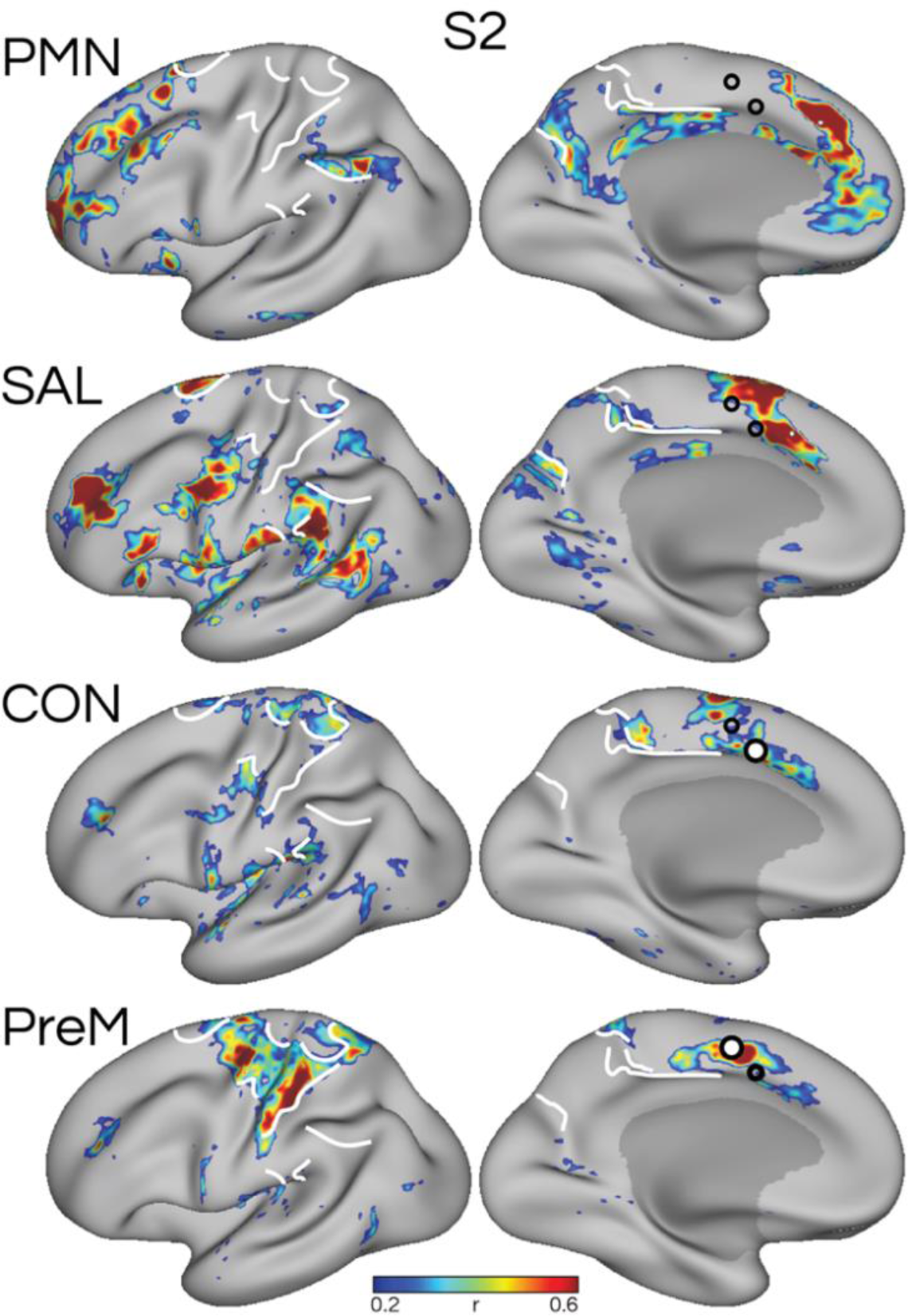
Targeted analysis confirms that the parietal memory network (PMN), salience network (SAL), and cingulo-opercular network (CON) are distinct, and are distinct from a premotor network (PreM) that surrounds primary somatomotor regions. Seeds were chosen from the dorsomedial prefrontal cortex in an anterior-posterior progression to target PMN, SAL, CON, and PreM in S2 (two additional subjects (S3 and S7) are shown in Fig. 6.) that provided particularly good separation between the networks in the high-resolution dataset. This analysis showed that SAL and CON are distinguishable from each other, and the premotor network, with each network occupying distinct regions of the cortex. White lines serve as hand-drawn landmarks for comparing across panels. White circles indicate the seed used to define the network shown in that panel, and black hollow circles represent seeds for the other networks shown.

**Supplementary Figure S19:**
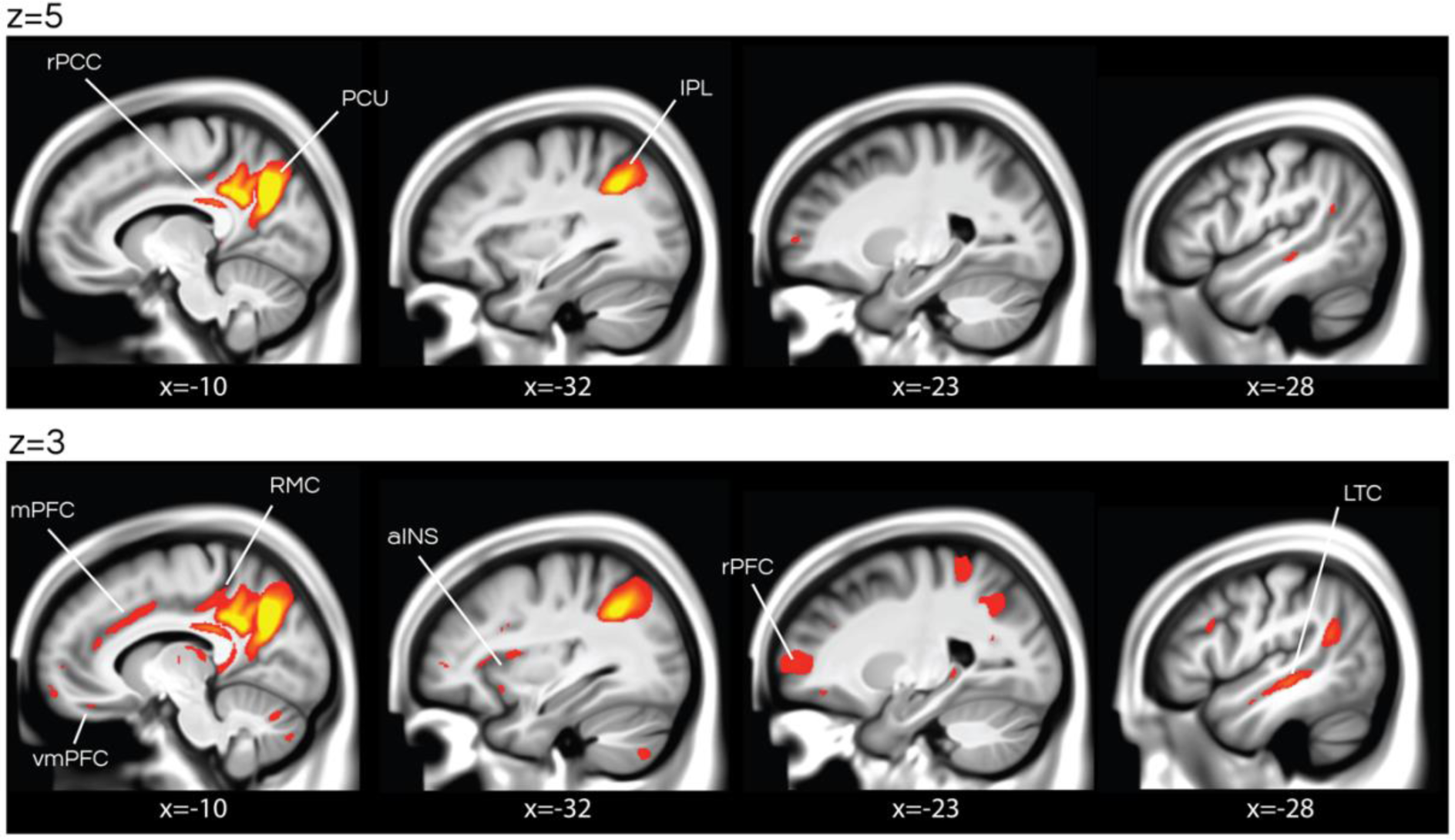
Group-averaged resting-state functional connectivity maps from the UK Biobank recapitulate that the parietal memory network (PMN) is distributed when examined at lower thresholds. Sagittal views show “component 21” from an independent component analysis (ICA) at 25-dimensions of data from 4,181 UK Biobank participants, taken from (Miller et al., 2016). At the default visualization threshold (z = 5), the network exhibits the characteristic features of the PMN, with prominent regions in the posteromedial cortex and inferior parietal lobule and hints of the presence of smaller regions in the rPFC, RMC and LTC. At a lower threshold (z = 3), the network additionally includes regions distributed across the same regions we observed in individuals (Fig. 1; dPFC not shown). In addition, evidence for a lateral temporal region was observed. rPFC; rostral prefrontal cortex, aINS; anterior insula, IPL; inferior parietal lobule, PCU; precuneus, rPCC: rostral posterior cingulate cortex, RMC; ramus marginalis of the cingulate sulcus, mPFC; medial prefrontal cortex, vmPFC; ventromedial prefrontal cortex, LTC; lateral temporal cortex.

**Supplementary Table S1.** Large amounts of high-quality resting-state data were included for each participant, and divided into discovery, replication and triplication datasets. Resting-state runs obtained using high-field 7T high-resolution functional MRI as part of the Natural Scenes Dataset (Allen et al., 2021) were divided into a discovery dataset and 1-2 replication datasets. The table presents the number of good quality runs, a total data time used in analyses, and quality control metrics for the discovery and validation datasets of six individuals. The quality metrics include signal-to-noise ratio (tSNR), maximum absolute head motion (max motion), and maximum framewise displacement (max FD). The mean values are presented, with standard deviations in parentheses. Runs that fell below our quality control criteria (i.e., tSNR below 20, max motion greater than 2.0 mm, or max FD greater than 0.4 mm) were excluded from the full dataset, leading to the complete exclusion of 2 subjects.

|  | S1 | S2 | S3 | S4 | S6 | S7 |
| --- | --- | --- | --- | --- | --- | --- |
| *Discovery dataset* |  |  |  |  |  |  |
| # of runs | 17 | 6 | 8 | 6 | 7 | 6 |
| amount of data (min) | 85 | 30 | 40 | 30 | 35 | 30 |
| tSNR | 165.14 (38.39) | 134.52 (53.72) | 157.24 (36.48) | 141.65 (26.54) | 99.19 (23.36) | 157.58 (37.79) |
| max motion (mm) | 0.43 (0.11) | 0.74 (0.26) | 0.48 (0.17) | 0.67 (0.30) | 0.68 (0.16) | 0.54 (0.18) |
| max FD (mm) | 0.26 (0.06) | 0.38 (0.05) | 0.26 (0.07) | 0.22 (0.09) | 0.18 (0.05) | 0.15 (0.04) |
| *Replication dataset* |  |  |  |  |  |  |
| # of runs | 9 |  | 8 | 6 | 6 | 6 |
| amount of data (min) | 45 |  | 40 | 30 | 30 | 30 |
| tSNR | 147.71 (33.03) | — | 159.79 (51.20) | 153.77 (35.57) | 108.33 (14.36) | 178.97 (45.88) |
| max motion | 0.36 (0.06) |  | 0.59 (0.23) | 0.62 (0.33) | 0.64 (0.30) | 0.61 (0.22) |
| max FD | 0.22 (0.03) |  | 0.28 (0.04) | 0.19 (0.08) | 0.14 (0.04) | 0.15 (0.03) |
| *Triplication dataset* |  |  |  |  |  |  |
| # of runs | 9 |  |  |  | 6 | 6 |
| amount of data (min) | 45 |  |  |  | 30 | 30 |
| tSNR | 148.94 (27.47) | — | — | — | 123.52 (35.30) | 133.95 (33.50) |
| max motion | 0.39 (0.14) |  |  |  | 0.68 (0.25) | 0.56 (0.21) |
| max FD | 0.24 (0.05) |  |  |  | 0.14 (0.07) | 0.15 (0.05) |

**Supplementary Table S2.**
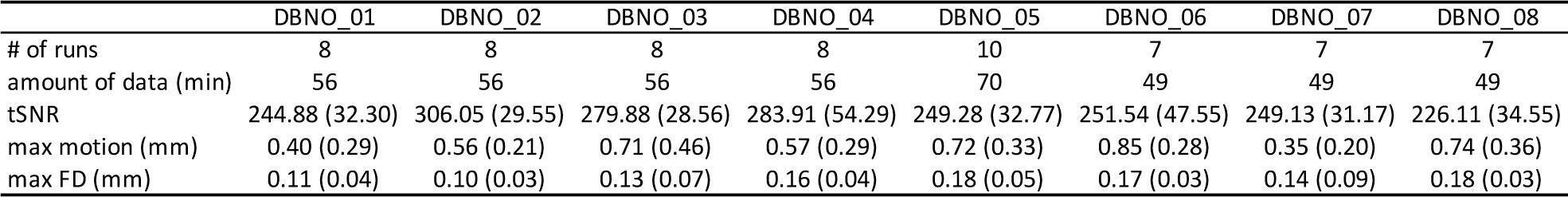
Data and quality control metrics for an independent 3T validation dataset that included eight participants. The table presents the number of runs included, total duration of data included in analyses, and quality control metrics including signal-to-noise ratio (tSNR), maximum absolute head motion (max motion), and maximum framewise displacement (max FD). The mean values are shown, with standard deviations in parentheses.

|  | DBNO_01 | DBNO_02 | DBNO_03 | DBNO_04 | DBNO_05 | DBNO_06 | DBNO_07 | DBNO_08 |
| --- | --- | --- | --- | --- | --- | --- | --- | --- |
| # of runs | 8 | 8 | 8 | 8 | 10 | 7 | 7 | 7 |
| amount of data (min) | 56 | 56 | 56 | 56 | 70 | 49 | 49 | 49 |
| tSNR | 244.88 (32.30) | 306.05 (29.55) | 279.88 (28.56) | 283.91 (54.29) | 249.28 (32.77) | 251.54 (47.55) | 249.13 (31.17) | 226.11 (34.55) |
| max motion (mm) | 0.40 (0.29) | 0.56 (0.21) | 0.71 (0.46) | 0.57 (0.29) | 0.72 (0.33) | 0.85 (0.28) | 0.35 (0.20) | 0.74 (0.36) |
| max FD (mm) | 0.11 (0.04) | 0.10 (0.03) | 0.13 (0.07) | 0.16 (0.04) | 0.18 (0.05) | 0.17 (0.03) | 0.14 (0.09) | 0.18 (0.03) |

